# Integrating a tailored recurrent neural network with Bayesian experimental design to optimize microbial community functions

**DOI:** 10.1101/2022.11.12.516271

**Authors:** Jaron C. Thompson, Victor M. Zavala, Ophelia S. Venturelli

## Abstract

Microbiomes interact dynamically with their environment to perform exploitable functions such as production of valuable metabolites and degradation of toxic metabolites for a wide range of applications in human health, agriculture, and environmental cleanup. Developing computational models to predict the key bacterial species and environmental factors to build and optimize such functions are crucial to accelerate microbial community engineering. However, there is an unknown web of interactions that determine the highly complex and dynamic behaviors of these systems, which precludes the development of models based on known mechanisms. By contrast, entirely data-driven machine learning models can produce physically unrealistic predictions and often require significant amounts of experimental data to learn system behavior. We develop a physically constrained recurrent neural network that preserves model flexibility but is constrained to produce physically consistent predictions and show that it outperforms existing machine learning methods in the prediction of experimentally measured species abundance and metabolite concentrations. Further, we present an experimental design algorithm to select a set of experimental conditions that simultaneously maximize the expected gain in information and target microbial community functions. Using a bioreactor case study, we demonstrate how the proposed framework can be used to efficiently navigate a large design space to identify optimal operating conditions. The proposed methodology offers a flexible machine learning approach specifically tailored to optimize microbiome target functions through the sequential design of informative experiments that seek to explore and exploit community functions.

**Author summary:** The functions performed by microbiomes hold tremendous promise to address grand challenges facing society ranging from improving human health to promoting plant growth. To design their properties, flexible computational models that can predict the temporally changing behaviors of microbiomes in response to key environmental parameters are needed. When considering bottom-up design of microbiomes, the number of possible communities grows exponentially with the number of organisms and environmental factors, which makes it challenging to navigate the microbiome function landscape. To overcome these challenges, we present a physically constrained machine learning model for microbiomes and a Bayesian experimental design framework to efficiently navigate the space of possible communities and environmental factors.

## 2 Introduction

Microbial communities have the potential to perform a variety of functions, including the ability to convert carbon-rich waste products into valuable compounds[1, 2], perform biological nitrogen fixation to improve agricultural yields[3], detoxify waste from the environment [4], and modulate vertebrate host phenotypes [5]. However, designing microbial communities from the bottom-up to perform desired functions is a major challenge due to unknown mechanisms of interaction, limited ability to observe and quantify all aspects of such systems (e.g. metabolites utilized and produced by constituent community members). Further, the design space of species and environmental factors for optimizing a microbiome target function is large and difficult to systematically navigate. Developing models that predict the temporal behaviors of communities from data and identify environmental conditions and combinations of species predicted to have optimized functions has emerged as a promising avenue to direct microbiome engineering[6].

Since microbiomes have large design spaces, high-throughput experiments coupled to computational modeling can be powerful for understanding and engineering microbial communities from the bottom-up[7, 8, 9]. Mathematical models that predict system behavior have become essential tools to understand complex biological processes [10], and recent studies have successfully applied a model guided approach to understand and optimize microbial community functions [9, 5]. Developing models of the microbiome from first principles is difficult due to unknown interactions as well as a limited understanding of the mechanisms that underlie these interactions [11]. Machine learning methods that can learn how microbial species interact in different environments from experimental data are thus compelling approaches to address this limitation. Neural networks are flexible machine learning models that can predict complex behavior for a broad class of systems [12]. Recurrent neural networks (RNNs), in particular, are powerful neural network architectures that can exploit multivariate time series data to learn dynamic behaviors [13]. For example, Baranwal et al. [14] showed that RNNs could model microbial community dynamics with greater accuracy than standard ecological models that are confined by a strict set of assumptions, such as the generalized Lotka-Volterra (gLV) model. In addition to improved prediction performance of species growth dynamics, the model was able to accurately forecast the production of health relevant metabolites given an initial profile of species abundances (i.e. species presence or absence). In addition, an RNN model trained on time series measurements of human gut microbiome composition data tailored for classification of food allergy achieved the best prediction accuracy compared to other machine learning methods [15].

While highly flexible, key limitations of applying machine learning models such as RNNs to physical systems is that they can produce unrealistic predictions (e.g. negative species abundances) and that they can require significant amounts of experimental data for training. Machine learning models are capable of making unrealistic predictions when the training data set (i.e. data used to build the model) is insufficient to constrain the model to match system behavior. When some mechanistic insights are known, embedding physical constraints into machine learning models can reduce the amount of data required for training and can result in improved prediction performance [16, 17]. Physically constrained machine learning models are especially promising for modeling biological systems [18] because these constraints can potentially improve a model’s ability to extrapolate beyond the regime explored in the training set despite limited or noisy data [19, 20]. In the computational biology field, for instance, neural networks have been used in concert with mechanistic ordinary differential equation (ODE) models to infer complex non-linear dynamics of partially observable biological systems [21]. In addition to incorporating physical constraints, experimental design strategies that optimize the information content of experimental data can reduce the amount of data needed to train a predictive model.

The collection of data used to inform machine learning models requires taking measurements of system properties, which is often time-consuming and expensive. Consequently, the selection of an informative set of experiments is crucial for developing models that capture system properties, while minimizing time and resources spent on performing experiments [22]. To achieve this goal, determining an optimal set of experiments that minimizes either model prediction uncertainty or uncertainty in parameter estimates has been widely used to optimize the information content of experiments for studying biological systems [23, 24, 25, 26]. Bayesian experimental design naturally integrates previously observed data to inform the selection of new experimental conditions. This enables a sequential strategy that uses all previously collected data to inform future iterations of model fitting, experimental design, and data acquisition. These approaches use acquisition functions that aim to quantify information content and predict system performance under potential sets of experiments. A widely used acquisition function is called the expected information gain (EIG), which quantifies how well an experimental design is expected to constrain estimates of model parameters [27, 26, 28]. While the EIG provides a principled acquisition function to design new experimental conditions, it is typically intractable to compute analytically for non-linear models and is slow to evaluate even when approximate approaches are used.

While most applications of Bayesian experimental design have focused on conducting experiments to refine a model, experimental design strategies have rarely been used in the field of systems biology for the purpose of seeking conditions that optimize properties of the system (e.g. production of a valuable compound or pathogen inhibition) for target applications. However, Bayesian experimental design can be tailored to provide a powerful goal-oriented framework that can leverage a flexible class of models to propose experimental conditions that have the dual objective of mitigating model uncertainty and optimizing system performance [27, 29]. For example, Bayesian optimization is an experimental design technique whose purpose is to efficiently optimize system properties and has been used in many fields ranging from synthetic biology [30] to aerospace engineering [31]. Bayesian optimization typically uses a Gaussian process to model system performance directly from experimental data. While Gaussian process models provide a natural and computationally-tractable approach to construct acquisitions functions [32], they cannot easily model the dynamic behavior of multivariate systems [33]. Another widely used goal-oriented experimental design strategy is called response surface methodology, which proposes experiments to build a performance function that is then optimized to find the best operating conditions. However, this approach is typically limited to linear models that use certain types of performance functions [34]. In addition, updated experimental designs need to be defined manually, as opposed to selected by the model.

We address gaps in model guided experimental design of microbial communities by developing and applying a physically constrained RNN architecture tailored to predict microbial community dynamics and target functions (e.g. production of specific metabolites) in response to environmental inputs. The proposed model outperforms other representative machine learning methods in the prediction of species abundances and metabolite concentrations using experimental data composed of unique human gut communities (> 10 species). Equipped with this model, we present an experimental design framework to optimize microbial community functions that leverages an information theoretic approach to select a set of experimental conditions that collectively exploit system functions and fill knowledge gaps in the model. We demonstrate the capability of the overall framework to minimize the number of experiments necessary to identify optimal operating conditions that maximize production of a desired metabolite using a mechanistic multi-species microbial community model. In sum, our framework uniquely integrates sequential Bayesian experimental design with a RNN tailored to predict and optimize dynamic microbial community behaviors.

## 3 Results

### 3.1 Design of microbial communities using a physically constrained recurrent neural network

Machine learning models can generate physically unrealistic predictions for physical systems. To address this limitation, we present the Microbiome Recurrent Neural Network (MiRNN), a modified RNN architecture that eliminates the possibility of predicting physically unrealistic species abundances and metabolite concentrations. We leverage a Bayesian inference method for parameter estimation, hyper-parameter optimization, and quantification of prediction uncertainty. A model-guided approach is used to identify a set of experimental conditions that collectively maximize information content of different experimental designs and design objectives. Our framework allows for the selection of an optimal set of experimental conditions that are collectively informative, as opposed to the selection of a single experimental condition, which can be applied to high-throughput experimental designs. The proposed methodology is illustrated in Figure 1. In the design phase, the MiRNN is combined with an acquisition function, *f*, to rank experimental designs based on predicted outcomes and the expected information gain (EIG) from a space of all possible experimental conditions, denoted as 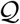. The acquisition function is composed of two parts, one that quantifies the expected profit of an experimental design and one that quantifies the information content of an experimental design. The highest ranked design, 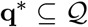, is tested to generate experimental data in the test phase. The resulting data is used to update the MiRNN model in the learn phase. The updated model is used to design the next experiment, completing the design, test, learn (DTL) cycle. DTL cycles can be repeated until convergence or until a desired objective is achieved.

**Figure 1:**
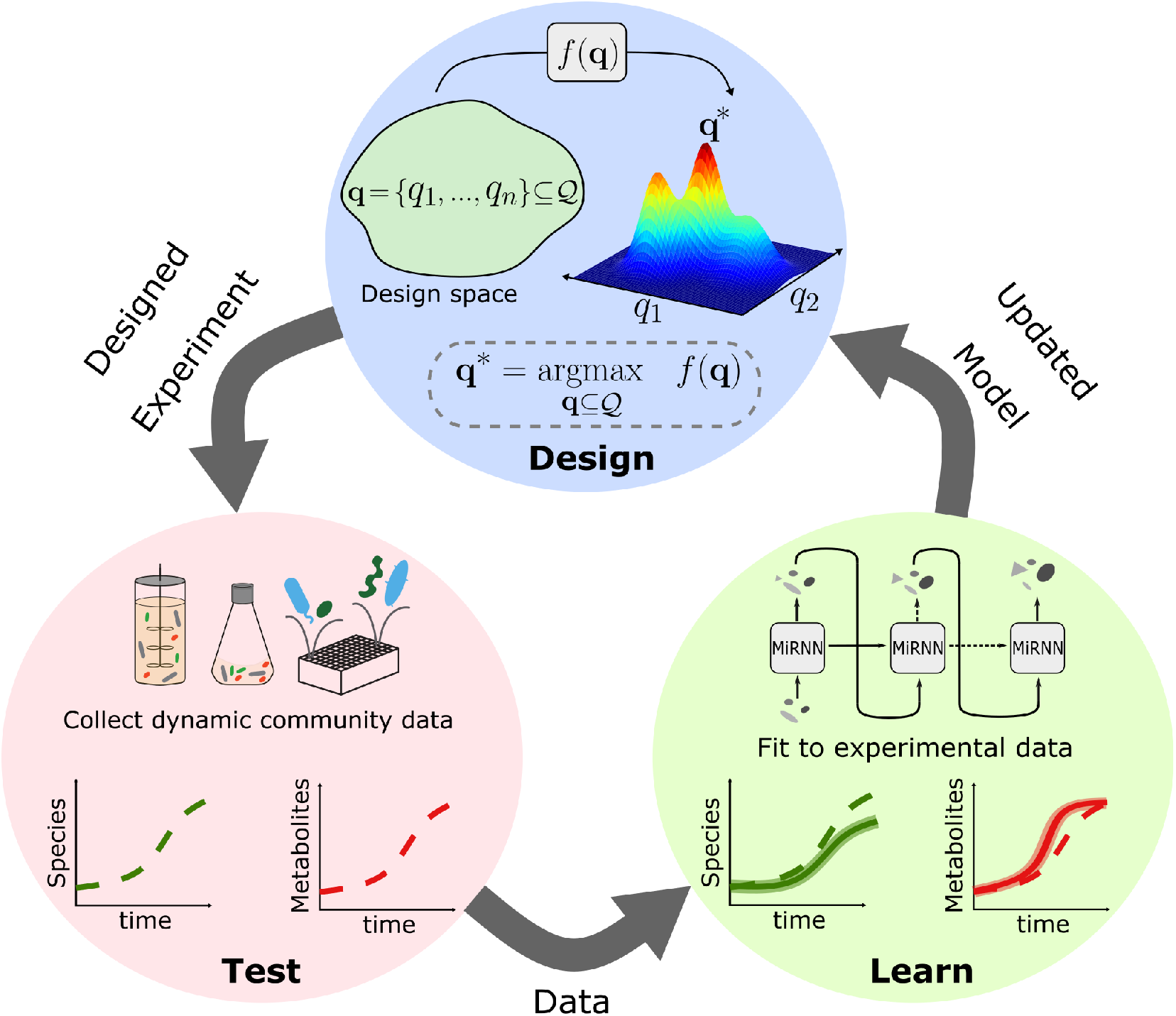
The Microbiome Recurrent Neural Network (MiRNN) learns system dynamics and proposes new designs. (**Design**) An experimental design space, denoted as 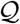, is a set of individual experimental conditions, **q**, where a particular condition could, for example, be a set of species in a community or the initial concentrations of resources. MiRNN predictions of outcomes for a set of experimental conditions, **q**, are evaluated by an acquisition function, *f*, which balances the expected information gain (EIG) of an experimental design and its expected profit to evaluate the optimality of experimental designs. (**Test**) The optimal experimental design, **q**\*, defines a set of experimental conditions to be observed experimentally. Measurements of these conditions are collected in the test phase. (**Learn**) Data collected in the test phase, and all previously collected data, are used to fit an updated MiRNN model. Once fit to the newly acquired data, the updated MiRNN model can be used again in the design phase to complete the design, test, learn cycle.

### 3.2 Comparison of the Microbiome Recurrent Neural Network (MiRNN) model to an unconstrained recurrent neural network

Microbiomes produce and degrade a myriad of metabolites, which mediate interactions with constituent community members and can be exploited to our benefit. To test the ability of the MiRNN to predict species abundance and metabolite concentration over time, we evaluated the model’s prediction performance on experimental data in which the absolute abundances of 25 diverse and prevalent human gut species and the concentrations of four major health-relevant metabolites (acetate, butyrate, lactate and succinate) were measured over time[9, 14] (Fig. 2a). In particular, butyrate produced by gut microbiota has beneficial effects on human health and disease, including promoting homeostasis in the colon [35, 36] and protecting against metabolic disorders [37]. The ability to predict metabolite concentrations such as butyrate as a function of the presence and absence of individual species could inform the design of next-generation defined bacterial therapeutics. To test our model’s predictive capability, we used an experimental data set consisting of 95 unique subsets of the 25 member community that were inoculated in equal species proportions *in vitro*. Species abundances and metabolite concentrations were measured every 16 hours for a total of 48 hours to characterize community assembly and metabolite dynamics.

**Figure 2:**
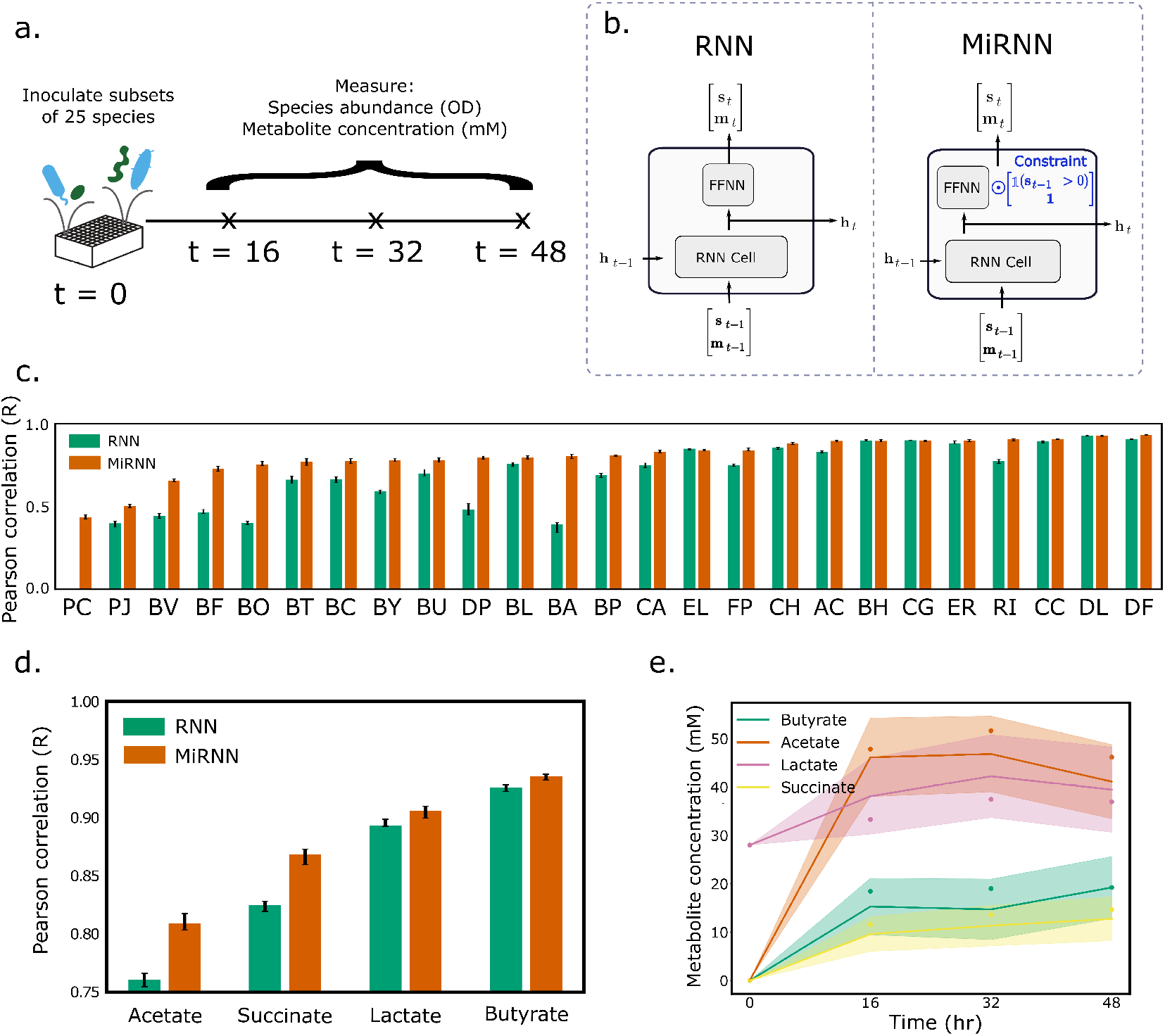
The predictive capability of the MiRNN outperforms unconstrained RNN model. (a.) Schematic of experiment in which 95 unique microbial consortia were selected from a set of 25 health relevant human gut bacteria. After inoculation, species abundances and metabolite concentrations were measured at 16 hour intervals over a course of 48 hours. (b.) A comparison of the MiRNN architecture to a standard RNN, where the constraint highlighted in blue prevents the model from predicting the spontaneous emergence of a species. (c.) Comparison RNN and MiRNN performance in species predictions according to the coefficient of determination between predictions and measured values. The height of the bars and error bars correspond to the median and interquartile range in prediction performance after running 20-fold cross-validation over 10 trials, where samples were randomly shuffled in each trial. (d.) Same as in panel (c.), except that metabolite prediction performance is shown. (e.) Representative temporal changes in MiRNN predicted metabolite concentrations, where measured values are shown as dots, the mean predicted value is shown as a line, and the uncertainty region shows ± 1 standard deviation.

We use a conventional RNN (left) and the MiRNN (right) to predict species abundance and metabolite concentrations at each time interval given an initial condition (Fig. 2b). The MiRNN is constrained such that species and metabolite concentrations cannot be negative and species absent from the given inoculum cannot be present at later time points. In contrast, an unconstrained model could predict negative species abundances or metabolite concentrations or spontaneous appearance of species that were not initially present (Fig. S1). To investigate differences in prediction performance of the MiRNN and RNN models, we performed 20-fold cross-validation by randomly partitioning the data into 20 unique sets of samples, training on 19 subsets, testing on the remaining subset, and then repeating for each combination of training and testing data so that all samples were subject to held-out testing. Because the partitioning of the data subsets is random, we repeated 20-fold cross-validation over 10 trials to evaluate the variation in prediction performance. On held-out data, predictions of species abundances using the MiRNN displayed a higher median Pearson correlation than the unconstrained RNN for 23 of the 25 species (Fig 2c), indicating that the incorporation of a physical constraint improved the model’s ability to predict species abundance. Although the constraint does not directly impact predictions of metabolite concentrations, the MiRNN outperformed the RNN in predictions of acetate and succinate. The MiRNN and RNN displayed similar prediction performance of lactate and butyrate, which displayed the highest prediction performance of the four metabolites (Fig. 2d,e). A representative trajectory shows the predicted distribution (mean **±**1 standard deviation) of each metabolite compared to measured values (Fig. 2e).

We compared the prediction performance of the MiRNN and RNN models to a Long-Short-Term-Memory (LSTM) model developed by Baranwal et al. [14] that was shown to accurately predict community dynamics and metabolite profiles. Similar to the MiRNN and RNN, the flexibility of the LSTM is governed by the dimension of the hidden layer, which was chosen to have 4096 hidden units in order to predict both species abundances and metabolite concentrations. For all analyses in this study, the hidden layer in the MiRNN and RNN contained 16 hidden units. Consequently, the LSTM proposed by Baranwal et al. contains orders of magnitude more parameters (*n_θ_* = 67, 735, 581) than the MiRNN and RNN (*n_θ_* = 1, 245), and thus offers greater flexibility. However, the large number of model parameters in the LSTM model makes it challenging to perform the Bayesian parameter inference method used to train the MiRNN and RNN models [38], since this requires the computationally expensive task of inverting a matrix with dimension equal to the square of the number of parameters [12]. Although the LSTM significantly outperformed the RNN when comparing to the median prediction performance of species (Fig. S2a) and metabolites (Fig. S2b) (paired t-test *p* < 1 × 10^-3^, *n* = 29), the deficit in prediction performance of the RNN was recovered by the MiRNN (Fig. S2c). In sum, the MiRNN prevents physically unrealistic predictions and enables the use of a model with substantially fewer parameters without sacrificing prediction performance. Reducing the number of parameters makes Bayesian inference methods more tractable, which provides a systematic method to determine model prediction uncertainty and optimize experimental designs.

We evaluated the quality of MiRNN prediction uncertainty on held-out data, since the ability to identify poorly understood conditions is crucial for selecting informative experimental designs that aim to fill knowledge gaps in the model. Evaluation of the log-likelihood of held-out testing data is a widely used approach to demonstrate a model’s ability to use prediction uncertainty to capture the variation in prediction error [39, 40]. Briefly, the log-likelihood (Eq. 5.3) of held-out data will be higher when model predicted uncertainty is small for predictions that are close to measured values and when model prediction uncertainty is large for predictions that are further away from measured values (Fig. S3a,b). We compared the log-likelihood of held-out data using a null model where the uncertainty in each prediction was estimated using a fixed variance, **Σ**_*y*_, to the log-likelihood using the condition-dependent model predicted variance given by Eq. 5.9.

The fixed estimate of **Σ**_*y*_ was computed using the expectation-maximization algorithm (SI appendix), which reflects the covariance in model prediction error on the training data. In this sense, **Σ**_*y*_ is the best guess of the variance that can be attributed to measurement noise. Uncertainty due to measurement noise cannot be reduced by collecting more data and is sometimes referred to as aleatory uncertainty, while uncertainty that could be minimized by collecting more data is referred to as epistemic uncertainty [41], both of which are captured by the model predicted uncertainty. The predicted uncertainty therefore reflects the degree of uncertainty associated with each experimental condition (e.g. the model could have varying levels of certainty about metabolite concentrations based on information from different consortia of microbial species). For the 10 randomized k-fold trials, the log-likelihood of held-out data using predicted variance was, on average, greater than the log-likelihood using the fixed variance (Fig. S3c). The increase in the log-likelihood using model predicted uncertainty suggests that accounting for both aleatory and epistemic uncertainty improved estimates of the distribution of model predictions compared to a model that only accounted for aleatory uncertainty. The ability to assign greater uncertainty to less understood experimental conditions is a key attribute that enables efficient exploration of a high-dimensional experimental design space.

### 3.3 Optimization of the production of a key metabolite by a microbial community in a bioreactor

Mixed microbial communities cultured in bioreactors have many bioprocessing applications, including valorization of agricultural waste[42], production of medium chain fatty acids from carbon rich waste streams [1], and production of bioplastics as an alternative to petroleum-based plastics [43, 44].

Optimizing these functions requires manipulation of process control variables such as substrate feed rates, feed composition, pH, and gas exchange [45]. Although most bioprocessing applications have involved single organisms, microbial consortia have several advantages. These advantages include the ability to transform a wider range of available nutrients into valuable compounds by exploiting different metabolic niches and division of labor [46, 47] and robustness of target functions to environmental perturbations such as invasion [48, 49, 50].

Resources (i.e. nutrients) are key control knobs for manipulating microbial community metabolism. Therefore, we consider selection of different combinations of resources and the rate at which the feed containing these resources is added to a fed-batch bioreactor containing a 5-member microbial community to maximize the production of a valuable metabolite (e.g. medium or long chain fatty acid [1]). Although our modeling framework is generally applicable to other reactor operation modes, such as continuous culture, we chose to study fed-batch operation to highlight the model’s ability to capture strong time-dependent changes in resources and biomass. Fed-batch operation involves a feed of substrates to the reactor without any discharge from the reactor. This in turn yields time-dependent variation in reactor volume, cell density, and product concentrations. This example demonstrates the ability of the MiRNN to optimize a multidimensional system with respect to control inputs that are both static (selection of resources) and dynamic (selection of the feed rate). As the ground truth system, we simulate a modified consumer-resource model[51] embedded in a fed-batch bioreactor model that assumes growth-associated kinetics for metabolite production (i.e. the rate of metabolite production is proportional to species growth rate) [52]. Species interactions are governed by competition for a limited set of resources. The governing equations of the ground-truth model are

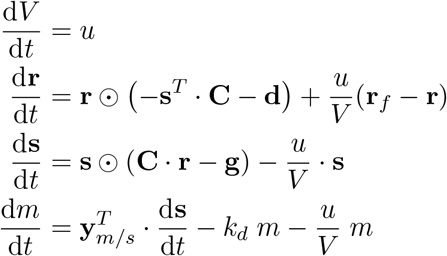

where **r** is a vector of resource concentrations in the reactor, **r**_*f*_ is a vector of resource concentrations in the feed, **s** is a vector of species that utilize a subset of the resources, **d** is a vector of resource degradation rates, **g** is a vector of minimum growth rates needed for each species to survive, *m* is the metabolite concentration, *y_m/s_* is a vector of yield coefficients, *k_d_* is the metabolite degradation rate, [**C**]_*ij*_ is the rate species *i* consumes resource *j*, and *u*(*t*) represents the rate at which the feed is added to the bioreactor. Details on the specification of ground truth model parameters are provided in the SI appendix. Due to competition for limiting resources, introducing different combinations of resources will determine the temporal changes in species abundance. The objective for the MiRNN was to model species growth, **s**(*t*), and metabolite production, *m*(*t*), and identify the optimal combination of 7 resources, 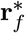, and the rate that the feed should be added over time, *u*\*(*t*), in order to maximize the total amount of target metabolite at the end of a 130 hour batch operation (Fig. 3a,b). We considered 20 possible feed profiles for each possible combination of resources, which resulted in 20 × (2^7^ – 1) = 2, 540 possible experimental conditions (SI appendix). This number of possible experimental conditions would not be feasible to exhaustively explore using generally low throughput bioreactor systems.

**Figure 3:**
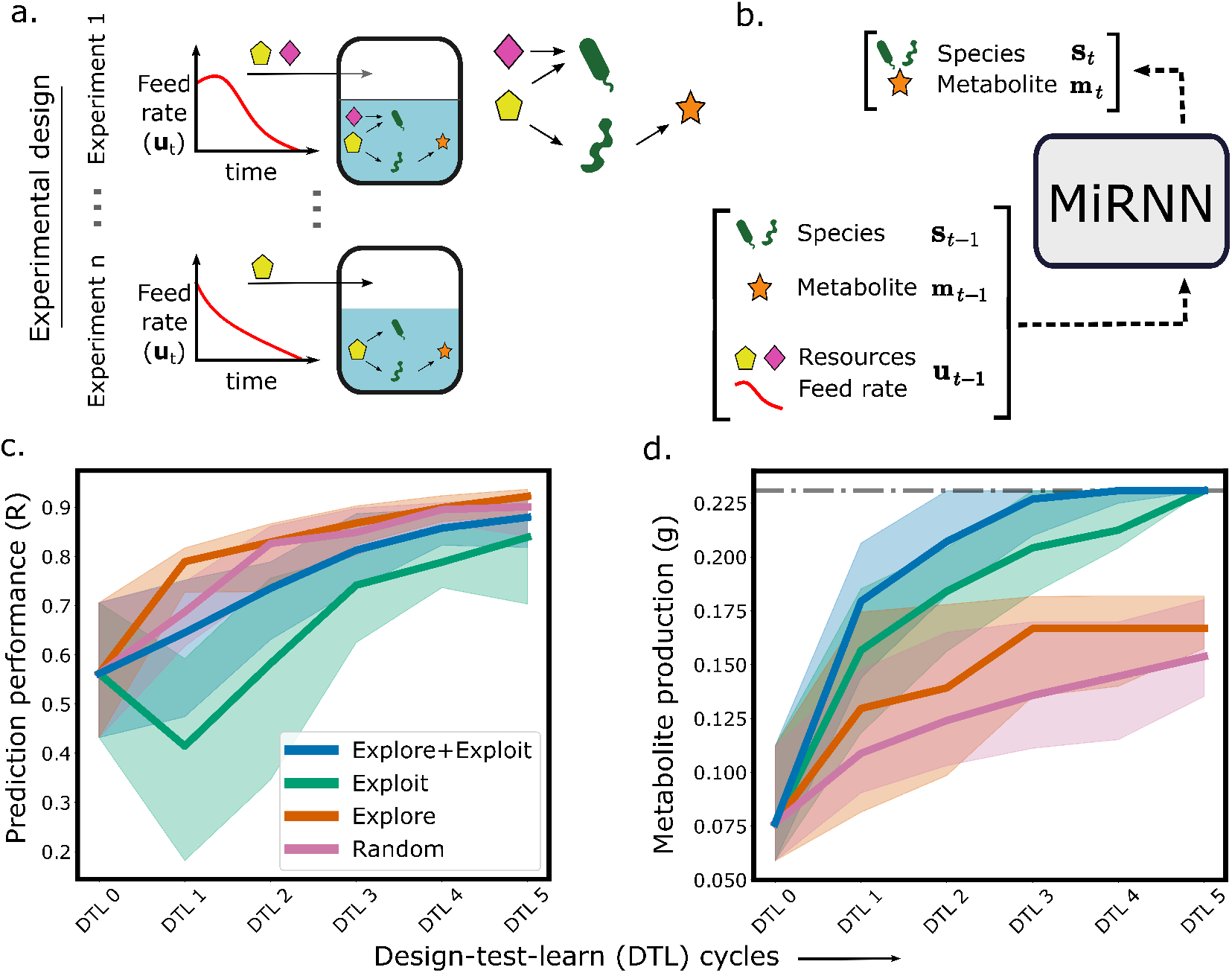
Optimization of resources and feed rate to maximize product. (a.) A schematic of the fed-batch bioreactor to be optimized, where the rate of a feed stream and the presence of resources in the feed (depicted as yellow and pink shapes) can both be adjusted in order to maximize production. Species (green shapes) that produce a valuable metabolite (orange star) compete for resources with species that do not produce the metabolite. (b.) A diagram that shows the inputs to the MiRNN model including species abundances, metabolite concentration, resource concentrations, and feed rate at time point *t* – 1. The model predicts species abundances and metabolite concentration at the next time step, *t*. Predicted species abundances and metabolite concentration are used as inputs to predict the next time step until the model has predicted the entire time course of the reaction. (c.) A comparison of prediction performance of end-point metabolite concentration between the proposed experimental design strategy that combines exploration and exploitation (blue) to pure exploitation (green), pure exploration (orange), and random sampling (purple). Solid lines show the median of the best recorded production (y-axis) up to each DTL cycle (x-axis) and uncertainty regions show the interquartile range computed over 30 trials each with random initial experimental designs. (d.) A comparison of metabolite maximization between proposed experimental design strategy that combines exploration and exploitation (blue) to pure exploitation (green), pure exploration (orange), and random sampling (purple). Solid lines show the median of the best recorded production (y-axis) up to each DTL cycle (x-axis) and uncertainty regions show the interquartile range computed over 30 trials each with random initial experimental designs.

We compared the effectiveness of four different experimental design strategies (random chance, pure exploration, pure exploitation, and exploration + exploitation) to find the bioreactor operating condition that maximized metabolite production. A pure exploration strategy seeks a set of experimental conditions that maximize the EIG, while the exploration + exploitation strategy evaluates experimental designs based on both the EIG and predicted outcomes. The variables subject to optimization included the resources in a feed stream and the rate at which the feed is added to a bioreactor over time (Fig. 3a, b). Starting with a randomly selected experimental design with five experimental conditions (DTL 0), each experimental design method was used to select the next set of five experimental conditions that would compose the next experimental design (DTL 1). Data collected from the next DTL cycle was used to update the model, which was then used to design the next DTL cycle, until five DTL cycles were completed. This process was repeated 30 times each with a different randomly selected set of five experimental conditions in DTL 0.

After each round of training, model predictions of end-point metabolite concentrations were compared to ground truth values for all 2, 540 experimental conditions to gauge how well the model learns system behavior (Fig. 3c). A pure exploration strategy results in the most accurate model performance after training on data from the first experimental design, while a pure exploitation strategy results in a decrease in model prediction performance due to sampling of redundant experimental conditions in a narrow region of the design space. The production levels from each experimental design strategy shows that all model guided approaches (exploitation+exploration, exploitation, and exploration) navigate to higher metabolite producing operating conditions than a random sampling strategy. The model guided experimental design strategy that combines exploitation and exploration outperforms pure exploitation (Fig. 3d), with significantly higher metabolite production in design cycles 1 (*p* = .0017), 2 (*p* = .0128), 3 (*p* = .0024), 4 (*p* = .0031), and 5 (*p* = .0203) according to a two-tailed paired t-test (n=30). The median prediction performance of end-point metabolite concentration of the exploitation+exploration strategy after training on all data collected up to DTL 2 was not the highest across different strategies (*R* = .735). Nevertheless, the median identified metabolite production in the next design cycle (DTL 3) was nearly optimal (.227 g). This implies that the model does not have to be highly accurate over the entire design space in order to be useful for seeking optimal conditions.

The ability of the MiRNN to predict both metabolite concentrations and species abundance over time can provide useful insight into the relationship between species abundances and system functions. This is in contrast to conventional Bayesian optimization approaches where a model (e.g. a Gaussian process) would be used to predict only metabolite production directly from the selection of resources and feed profile. To see how prediction of species abundances can provide insight, we can analyze MiRNN predictions of species abundances and metabolite production for the condition that was predicted to maximize metabolite production. The MiRNN predicted both high metabolite production and relatively high growth of species *s_2_* (Fig. S4d), suggesting that species *s_2_* produces the metabolite. This matches the ground truth model where the only species that produces the target metabolite is *s*_2_, 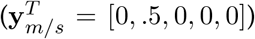. This agreement between model predictions and the ground truth system suggests that, when ground truth is not known, analyzing MiRNN predictions of system behavior under different experimental conditions can provide meaningful insights.

## 4 Discussion

Despite their potential, the bottom-up design of microbiomes remains a challenge due to the massive design space of possible microbial consortia and environmental inputs (e.g. resources). Further, mechanisms driving community behaviors are typically not known precluding the development of predictive computational models based on first principles. In this work, we present the Microbiome Recurrent Neural Network (MiRNN); a physically constrained RNN model tailored to predict the dynamics of species interactions from data and predict target community functions. We use an approximate Bayesian inference strategy to compute a posterior parameter distribution, which enables the quantification of model prediction uncertainty and the evaluation of the information content of potential experimental designs. The ability of the MiRNN to learn microbial community dynamics from previously acquired data and evaluate the information content of experimental designs enables a sequential design-test-learn strategy to efficiently seek experimental conditions that optimize community functions (Fig. 1).

Recent studies have underscored the need for an iterative design-test-learn strategy to build computational models that enable efficient exploration and exploitation of biological systems[53, 54], and in particular, microbial communities[55, 56]. Toward this end, we introduce the first physically constrained machine learning model to predict the dynamics of microbial communities and show that incorporating a physical constraint significantly improved the model’s ability to predict species abundances and metabolite concentrations using experimental data. Further, the model yielded comparable prediction performance to a previously developed LSTM model [14] despite more than a 50,000 fold reduction in the number of model parameters (Fig. S2). The reduction in model parameters makes the use of Bayesian inference techniques more tractable for the purpose of quantifying model prediction uncertainty and evaluating the information content of experimental designs. Model prediction uncertainty is used in active learning [57, 58], Bayesian optimization [32], and reinforcement learning [45]. The framework differs from most previous work on optimal experimental design of biological systems [23, 24, 26] since it leverages model uncertainty to select a set of experimental conditions for the purpose of optimizing a function of interest (exploitation and exploration), as opposed to designing experiments for the sole purpose of refining a model (exploration). Our results demonstrate that while a pure exploration strategy is the best approach for improving model predictions, it does not efficiently seek conditions that optimize a system objective. However, the proposed experimental design strategy that combines exploration with exploitation reduces the number of experiments needed to find optimal conditions compared to exploitation alone (Fig. 3c,d).

A limitation of the proposed methodology is that it relies on several approximations to enable fast selection of an experimental design, such as the assumption of Gaussian posterior parameter and predictive distributions. However, our model is an approximation of the ground truth system and despite imperfect predictions of system outcomes, the model is still able identify optimal experimental conditions (Fig. 3). We therefore expect an approximate estimate of the information content of experimental designs to be sufficient in most applications. However, determining the effectiveness of our proposed experimental design framework to optimize systems where conditional distributions are known to be non-Gaussian could be an area of future work. A limitation of the MiRNN, and any neural network based model, is that it offers limited interpretability for extracting new knowledge about the system. To tackle this problem, methods to extract meaning from a trained model such as Local Interpretable Model-agnostic Explanations (LIME)[59] have been used to derive relationships between variables in a similar modeling framework applied to microbial communities [14]. Alternatively, model predictions under different experimental conditions can be useful for gleaning mechanistic insights. For example, analyzing model predictions under the experimental condition that resulted in optimized metabolite production in our bioreactor case study correctly suggested that species *s*_2_ was responsible for producing the target metabolite (Fig. S4). Additionally, a limitation of discrete time models such as RNNs is that they require time series data to be sampled at consistent time intervals, which is often not the case in biological data sets where time series measurements are taken at different time resolutions. To overcome this limitation, the time interval can be included as an additional feature to the model or continuous time models such as neural ordinary differential equations [60] could be explored in future work.

Our framework enables the optimization of microbial community functions using a sequential Bayesian experimental design strategy. Our approach is capable of incorporating time dependent inputs (e.g. feed rate of a bioreactor) as potential controls to modulate system behavior. We note that although the constraint was incorporated for the purpose of modeling physically consistent bacterial growth, the same model could be applied to other chemical reaction networks that exhibit autocatalytic behaviors. For optimization of synthetic microbial communities, we envision future applications of this framework to include the selection of microbial species and resources (e.g. fibers) to accelerate the discovery of bacterial therapeutics that produce beneficial metabolites and display robustness, the selection of microbial species and environmental conditions to increase biological nitrogen fixation for enhancing plant growth, and the design of microbial consortia with improved productivity of valuable chemicals such as medium chain fatty acids and bioplastics.

## 5 Methods

### 5.1 Microbiome Recurrent Neural Network (MiRNN) Model

RNNs are flexible machine learning models that can be applied to learn complex dynamical models directly from multivariate time series data. In this work, we present the Microbiome Recurrent Neural Network (MiRNN), illustrated in Fig 4, which is a modified RNN that aims to learn and predict the dynamic behavior of microbial communities. A more detailed description of the model architecture is shown in Fig. S5. Specifically, the MiRNN architecture aims to learn dynamic trajectories for species abundances and metabolites given a set of potentially time dependent inputs and encodes constraints that prevent prediction of negative species abundance or metabolite concentration and prevents the spontaneous emergence of a species.

**Figure 4:**
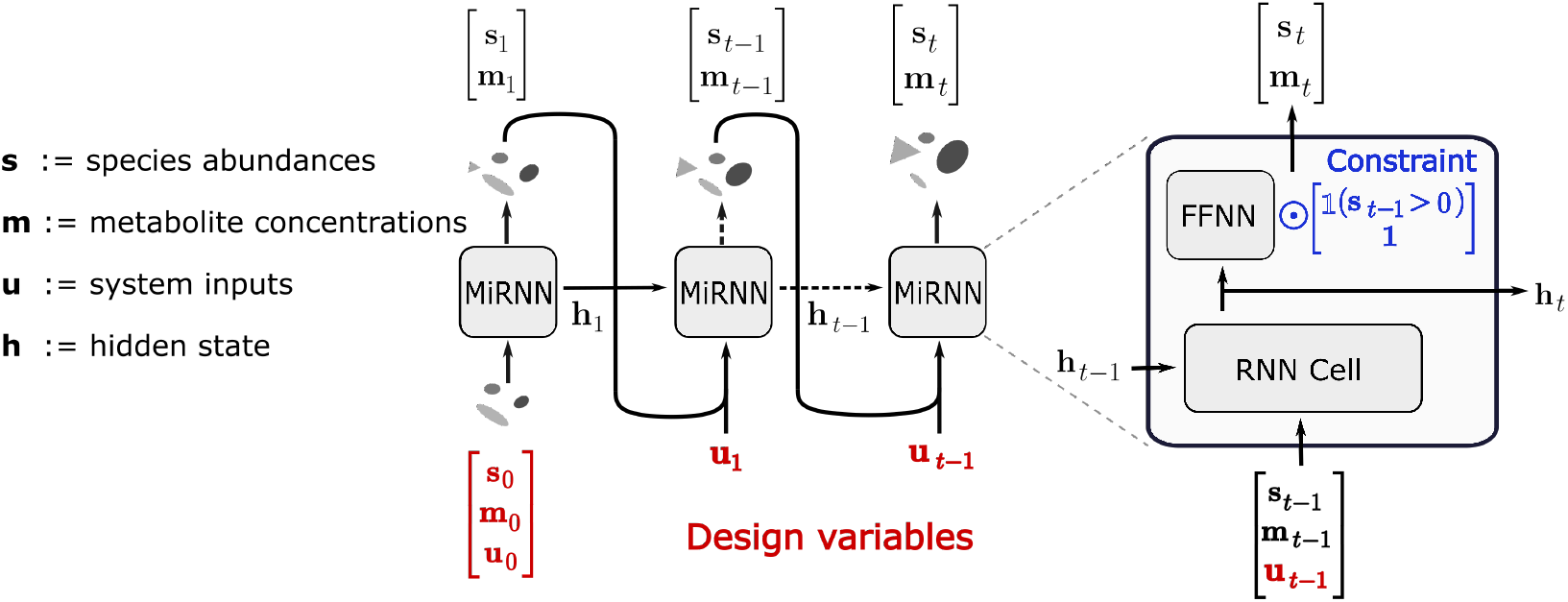
Microbiome Recurrent Neural Network architecture. Inputs to the RNN at time step *t* – 1 include the state of species abundances, metabolite concentrations, control inputs, and a latent vector that stores information from previous steps and whose dimension determines the flexibility of the model. The output from each MiRNN block is the predicted system state and the latent vector at the next time step, *t*. To avoid the physically unrealistic emergence of previously absent species, a constrained feed-forward neural network (FFNN) outputs zero valued species abundances if species abundances at the previous time step were zero.

We define a time horizon given by the index *t* = 0,…, *n_τ_*. The concentration of species at time *t* is denoted as 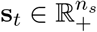. The concentration of metabolites at time *t* is denoted by 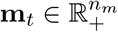. The value of the controls (inputs) at *t* is given by 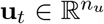. The dynamic evolution of the MiRNN model is given by a mapping of the form:

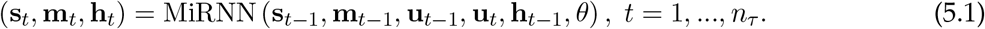

Here, 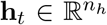 is a vector of latent variables at time *t* and 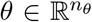 is a vector of model parameters. The latent variables propagate information from previous states in time. Increasing the dimension of the latent variable vector increases model complexity and flexibility, and can be selected using training data by maximizing the model evidence. For all analyses in this study, we set *n_h_* = 16. We note that the controls at step *t* – 1 and *t* are both fed into the model evolution to account for the possibility of encountering strong time dependent variations in the control. The outputs of the RNN are the predicted system state and the latent vector at step *t*.

The set of model parameters of the architecture is composed of the weights and biases *θ* = {**W**_*hh*_, **b**_*hh*_, **W**_*ih*_, **W**_*ho*_, **b**_*ho*_, **h**_0_}, which are learned from data. The MiRNN architecture is designed to prevent the physically unrealistic emergence of previously absent microbial species; this is done by introducing a logic block that sets the species abundances to zero if the abundances at the previous time step were zero. By applying the rectified linear unit activation function to model outputs, the state vector is strictly non-negative. Because the data are scaled to values between zero and one based on the maximum values in the training data, the ReLU activation eliminates the possibility of negative valued model outputs after applying the inverse scaling transformation.

In the context of experimental design, the control trajectories **u**_*t*_, *t* = 0,…, *n_τ_* and the initial states **s**_0_ and **m**_0_ are variables that we can manipulate to influence the evolution of the state trajectories, s_*t*_ (species) and **m**_*t*_ (metabolites) for *t* = 1,…, *n_τ_*. Observed (measured) variables are referred to as outputs or observables; here, we assume that species abundances and metabolite concentrations can be measured and we encapsulate the entire set of output variables in the vector 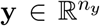. The manipulated variables are called design variables in an experimental design context. We refer to a particular choice of design variables as an experimental condition, which is denoted by the variable *q_i_*. We define an experimental design as a set of *n* experimental conditions, **q** = {*q*_1_,…,*q_n_*}. We denote the entire set of *m* ≥ *n* possible experimental conditions as 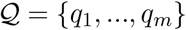.

### 5.2 Bayesian Estimation and Uncertainty Quantification

We use a Bayesian framework to estimate the parameters of the model from designed experiments and to quantify the uncertainty of the model predictions given such experiments. We assume that we have an initial experimental design **q** with associated experimental conditions *q_i_* indexed by *i* = 1,…,*n* each with corresponding observed outputs **y**(*q_i_*). The entire set of available data from an experimental design, **q**, is defined as the set 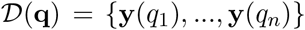. We assume that the output observations are contaminated by random noise as:

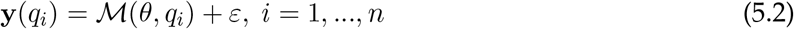

where 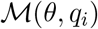 is the MiRNN output prediction at experimental condition *q_i_*, 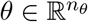 are the MiRNN parameters, and *ε* is a noise random variable with probability density 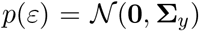. The matrix **Σ**_*y*_ describes the variance of the noise (this is a hyper-parameter that can be defined manually or can be inferred from data (SI appendix). We assume that the random noise over the multiple experiments is independent and identically distributed (i.i.d) and thus the model likelihood is given by:

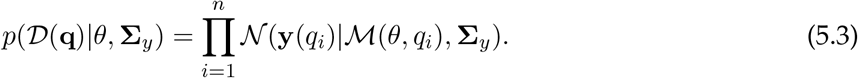

The prior over parameters is given by 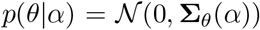, where 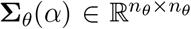 is the prior covariance (a diagonal matrix) and 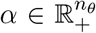 is a tunable hyper-parameter vector that inflates/deflates this covariance (SI appendix). From Bayes’ theorem, the posterior parameter distribution is proportional to the product of the likelihood and the prior. The mode of the posterior density provides the *maximum a posteriori* (MAP) estimate of model parameters and is obtained by minimizing the negative log of the posterior density with respect to *θ*,

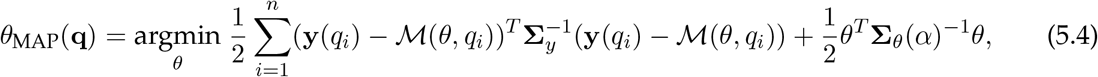

which we solve numerically using Newton’s method. The addition of the prior in the likelihood function encourages sparsity of the parameter estimates if *α* is made sufficiently large [61, 12].

In order to quantify the uncertainty of the parameter estimates, it is necessary to obtain their posterior density. Here, we use the so-called Laplace approximation; this assumes the posterior density is a Gaussian centered at *θ*_MAP_ (**q**) and with covariance given by the inverse of the Hessian matrix of the negative log posterior, which is approximated as:

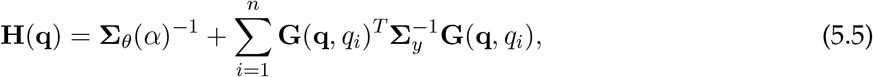

where 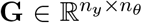 is a matrix of derivatives of the model with respect to its parameters (referred to as the sensitivity matrix) and given by:

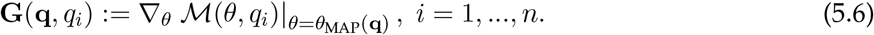

We note here that the Hessian matrix given by Eq. 5.5 is full rank due to the inclusion of the diagonal prior precision matrix. The posterior predictive distribution of the outputs for any experimental condition 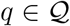 is found by marginalization over the posterior parameter distribution as:

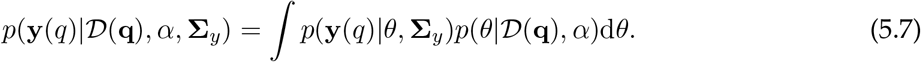

Obtaining an analytical expression for (5.7) requires linearization of the model prediction with respect to the parameters around *θ*_MAP_(**q**) to obtain a linear-Gaussian model [12],

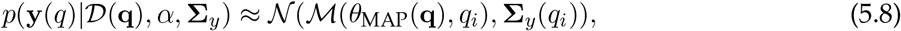

with

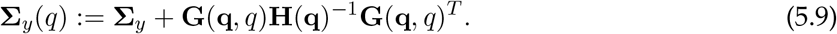

These expressions highlight how the design variables **q** propagate through the data 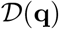, the calculation of the estimate *θ*_MAP_(**q**), and ultimately influence the uncertainty of the model predictions. As such, it is important to derive systematic procedures to determine such experiments.

### 5.3 Fast Bayesian Experimental Design to optimize information content and system outcomes

Bayesian experimental design methods use information from a previous experimental design, **q**^(*l*)^, to inform the selection of the next experimental design, **q**^(*l*+1)^. One commonly used strategy is to find **q**^(*l*+1)^ that maximizes the expected information gain (EIG), which is quantified by the expected Kullback-Leibler divergence between the parameter posterior and the current parameter distribution [27, 58],

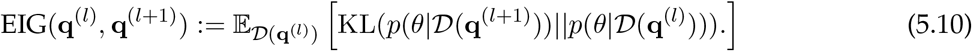

Using the model predictive distribution given by Eq. 5.8 and assuming that the posterior distribution is Gaussian, the EIG can be approximated as (SI appendix)

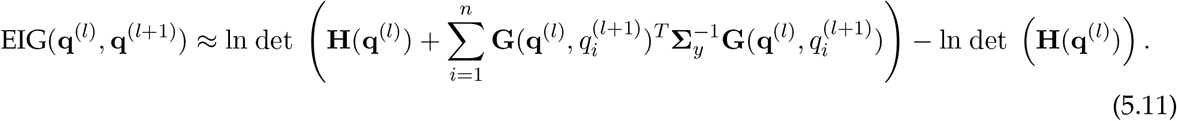

For linear-Gaussian models, experimental designs that maximize equation Eq. 5.11 are referred to as Bayesian D-optimal [27], because they maximize the determinant of the expected posterior precision matrix. Similarly, D-optimal experimental designs are often selected based on the determinant of the Fisher information matrix (FIM) [62], given by

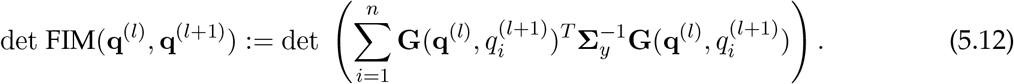

Although widely used in practice [63, 25, 64, 65], methods for experimental design based on maximizing either Eq. 5.11 or Eq. 5.12 can be computationally expensive since they require evaluating the determinant of a matrix with dimensions *n_θ_* × *n_θ_*. If the experimental design is composed of a single experimental condition, *q*^(*l*+1)^, it has been shown [27, 58] for linear-Gaussian models that the condition that maximizes the EIG is equivalent to the condition that maximizes the determinant of the prediction covariance due to the following identity (SI appendix),

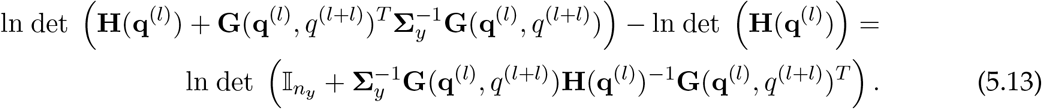

Since typically *n_y_* ≪ *n_θ_*, finding the experimental condition that maximizes prediction variance is a computationally efficient means of finding a Bayesian D-optimal condition; however, it is often desirable in experimental design applications to evaluate the information content of a set of *n* > 1 experimental conditions. We therefore present an expression that we show to be equivalent to Eq. 5.11 (SI appendix) and which generalizes Eq. 5.13 to compute the information content of *n* conditions,

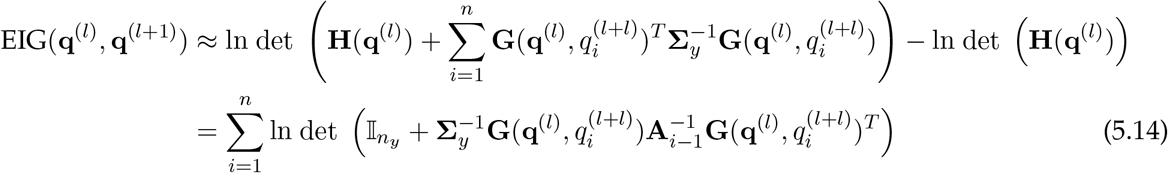

where

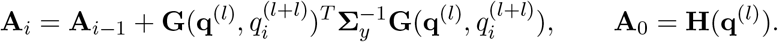

The matrix inverse in Eq. 5.14 can be efficiently computed using the Woodbury matrix identity,

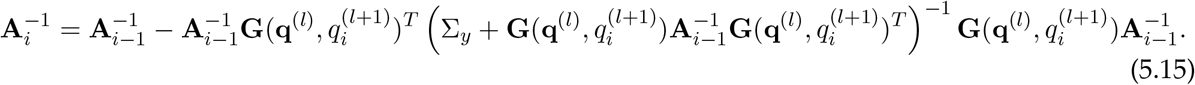

Using Eqs. 5.14 and 5.15, we can efficiently approximate the EIG of an experimental design with *n* experimental conditions by avoiding computing the determinant of a matrix whose dimension is *n_θ_* × *n_θ_* in favor of evaluating the determinant and inverse of n matrices each with dimension *n_y_* × *n_y_* (Fig. S6).

In this work, we aim to find experiments that maximize information content and optimize a profit function of interest. As such, we define an acquisition function that accounts for the predicted profit of experimental outcomes [27] as well as the expected information gain,

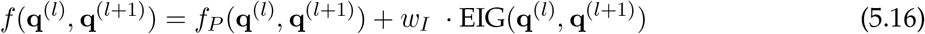

where *f_P_*(**q**^(*l*)^, **q**^(*l*+1)^) quantifies the predicted profit of the next design (e.g., total amount of product produced in each experiment). The profit function is an implicit function of the MiRNN model 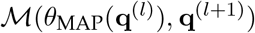 (i.e., the profit is predicted using the MiRNN model). The function EIG(**q**^(*l*)^, **q**^(*l*+1)^) quantifies the information content of the design **q**^(*l*+1)^) and is approximated using Eqs. 5.14 and 5.15. The parameter 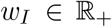 modifies the emphasis given to either profit or information content, and can be automatically adjusted to select for new experimental conditions as described in section 5.4. Given previously observed experimental designs **q**^(*l*)^ and a set 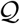 of all possible experimental conditions that could be tested, our goal is to select the next design 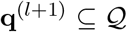 such that we maximize the acquisition function:

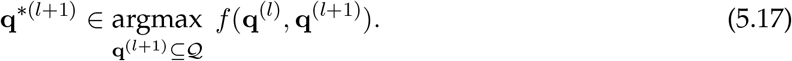

We note that the optimal experiments **q**\*^(*l*+1)^ are obtained based on best predicted performance (as predicted by the model); as such, these need to be tested in the real experimental system to obtain new outputs. This allows us to obtain a sequential experimental design framework in which we aim to progressively refine the model to maximize the profit function of interest.

#### Algorithm 1: Sequential Bayesian experimental design

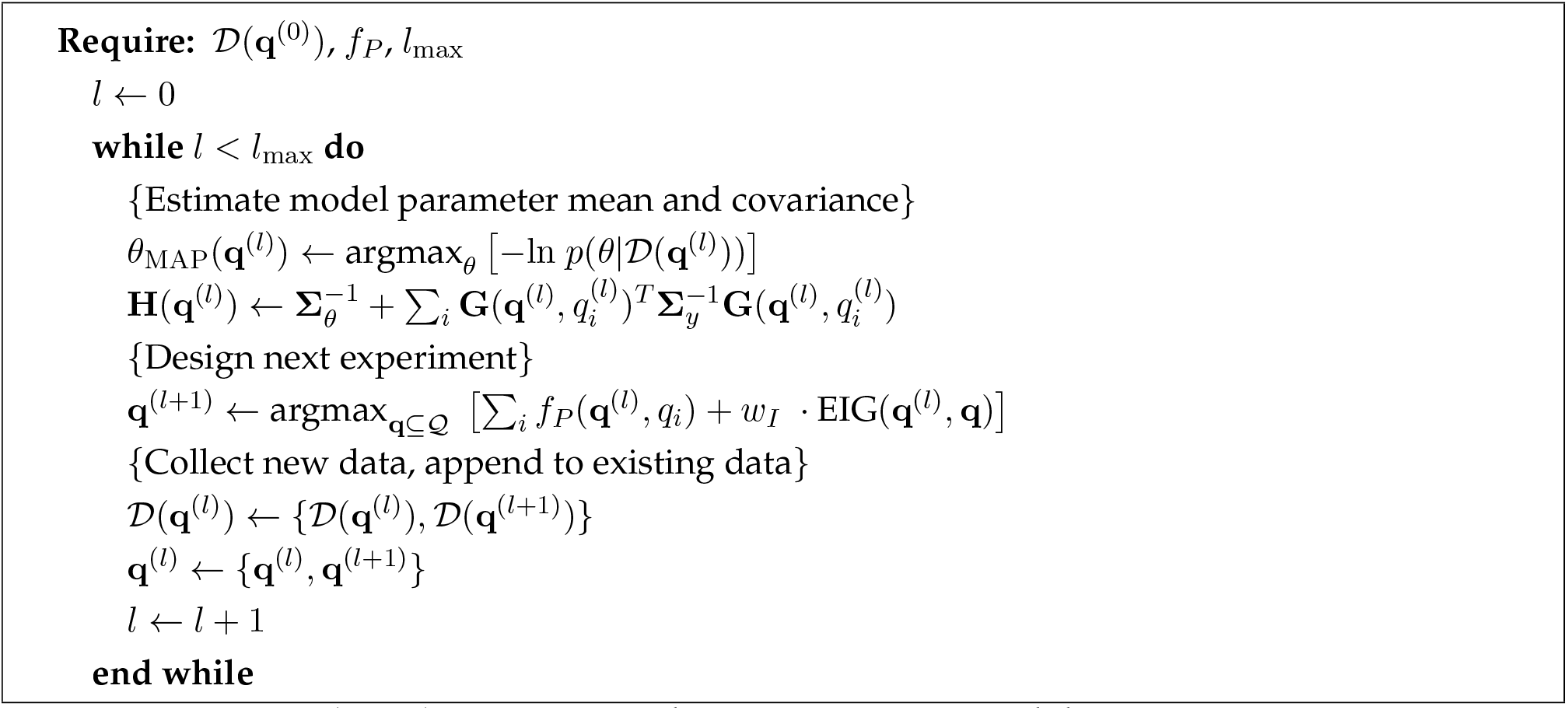

### 5.4 Greedy algorithm to search for optimal experimental designs

Finding the optimal next design **q**\*^(*l*+1)^ requires an exhaustive search over the set 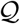 (particularly when the design variables are categorical). As expected, however, exhaustive enumeration would require evaluating *f*(**q**^(*l*)^, **q**^(*l*+1)^) for all 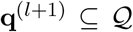 (which can be computationally prohibitive). As such, we implement a greedy search algorithm that works satisfactorily well in practice. It is important to emphasize that the experimental design framework has the final goal of maximizing the profit function (as opposed to just refine the model); as such, it searches for experiments in a more targeted manner and can improve profit without having a perfect prediction model. Greedy algorithms are often used as an approximate approach to optimize experimental designs [24]. Given a total number of conditions to include in the next design, *n*_q_(*l*+1), the search starts by finding an experimental condition that maximizes the profit function, 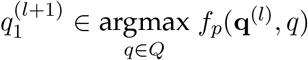. With **q**^(*l*+1)^ initialized as 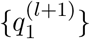, the *i* > 1 experimental condition is selected by determining

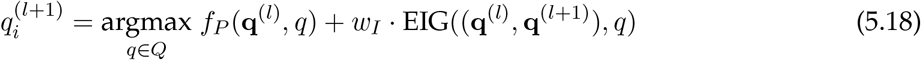

where *w_I_* is set to a small initial value (e.g. .0001) and gradually increased until 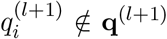. The process continues until a desired number of conditions are selected.

## 6 Author contributions

J.C.T., V.M.Z. and O.S.V. conceptualized research, wrote and edited the manuscript. J.C.T. wrote the original draft, wrote software, and performed research. O.S.V., J.C.T., and V.M.Z. analyzed and interpreted data. O.S.V. acquired funding.

## 8 Data availability

All code and data needed to reproduce the results have been shared as an installable Python package and documented Jupyter notebooks, which will be available at the time of publication from github.com/VenturelliLab.

## 9 SI appendix

The SI appendix provides additional methodological details on data pre-processing, evaluation of model prediction performance, hyper-parameter optimization, justification of the experimental design information function, fast evaluation of the experimental design information function, and generation of ground truth bioreactor model parameters.

## 10 Acknowledgements

We would like to thank Alfred Hero for his feedback in early stages of this project. This research was supported by funding from the Army Research Office (ARO) under grant number W911NF1910269, the National Institutes of Health under grant numbers R35GM124774 and R01EB030340.

## References

[1] Matthew J Scarborough, Griffin Lynch, Mitch Dickson, Mick McGee, Timothy J Donohue, and Daniel R Noguera. Increasing the economic value of lignocellulosic stillage through medium-chain fatty acid production. Biotechnology for biofuels, 11(1):1–17, 2018.

[2] Matthew T Agler, Catherine M Spirito, Joseph G Usack, Jeffrey J Werner, and Largus T Angenent. Chain elongation with reactor microbiomes: upgrading dilute ethanol to medium-chain carboxylates. Energy & Environmental Science, 5(8):8189–8192, 2012.

[3] Sanjana Kaul, Malvi Choudhary, Suruchi Gupta, and Manoj K Dhar. Engineering host microbiome for crop improvement and sustainable agriculture. Frontiers in Microbiology, 12:1125, 2021.

[4] Frank E Löffler and Elizabeth A Edwards. Harnessing microbial activities for environmental cleanup. Current Opinion in Biotechnology, 17(3):274–284, 2006.

[5] Richard R Stein, Takeshi Tanoue, Rose L Szabady, Shakti K Bhattarai, Bernat Olle, Jason M Norman, Wataru Suda, Kenshiro Oshima, Masahira Hattori, Georg K Gerber, et al. Computer-guided design of optimal microbial consortia for immune system modulation. Elife, 7:e30916, 2018.

[6] Christopher E Lawson. Retooling microbiome engineering for a sustainable future. Msystems, 6(4):e00925–21, 2021.

[7] Ophelia S Venturelli, Alex V Carr, Garth Fisher, Ryan H Hsu, Rebecca Lau, Benjamin P Bowen, Susan Hromada, Trent Northen, and Adam P Arkin. Deciphering microbial interactions in synthetic human gut microbiome communities. Molecular systems biology, 14(6):e8157, 2018.

[8] Susan Hromada, Yili Qian, Tyler B Jacobson, Ryan L Clark, Lauren Watson, Nasia Safdar, Daniel Amador-Noguez, and Ophelia S Venturelli. Negative interactions determine clostridioides difficile growth in synthetic human gut communities. Molecular systems biology, 17(10):e10355, 2021.

[9] Ryan L Clark, Bryce M Connors, David M Stevenson, Susan E Hromada, Joshua J Hamilton, Daniel Amador-Noguez, and Ophelia S Venturelli. Design of synthetic human gut microbiome assembly and butyrate production. Nature communications, 12(1):1–16, 2021.

[10] Ezio Bartocci and Pietro Lió. Computational modeling, formal analysis, and tools for systems biology. PLoS computational biology, 12(1):e1004591, 2016.

[11] Alicia Sanchez-Gorostiaga, Djordje Bajić, Melisa L Osborne, Juan F Poyatos, and Alvaro Sanchez. High-order interactions distort the functional landscape of microbial consortia. PLoS Biology, 17(12):e3000550, 2019.

[12] Christopher M Bishop and Nasser M Nasrabadi. Pattern recognition and machine learning. Springer, 2006.

[13] Ian Goodfellow, Yoshua Bengio, and Aaron Courville. Deep Learning. MIT Press, 2016. http://www.deeplearningbook.org.

[14] Mayank Baranwal, Ryan L Clark, Jaron Thompson, Zeyu Sun, Alfred O Hero, and Ophelia S Venturelli. Recurrent neural networks enable design of multifunctional synthetic human gut microbiome dynamics. eLife, 11:e73870, jun 2022.

[15] Ahmed A Metwally, Philip S Yu, Derek Reiman, Yang Dai, Patricia W Finn, and David L Perkins. Utilizing longitudinal microbiome taxonomic profiles to predict food allergy via long short-term memory networks. PLoS computational biology, 15(2):e1006693, 2019.

[16] Shubhendu Kumar Singh, Ruoyu Yang, Amir Behjat, Rahul Rai, Souma Chowdhury, and Ion Matei. Pi-lstm: Physics-infused long short-term memory network. In 2019 18th IEEE International Conference On Machine Learning And Applications (ICMLA), pages 34–41. IEEE, 2019.

[17] Jared Willard, Xiaowei Jia, Shaoming Xu, Michael Steinbach, and Vipin Kumar. Integrating scientific knowledge with machine learning for engineering and environmental systems. ACM Computing Surveys (CSUR), 2021.

[18] Grace CY Peng, Mark Alber, Adrian Buganza Tepole, William R Cannon, Suvranu De, Savador Dura-Bernal, Krishna Garikipati, George Karniadakis, William W Lytton, Paris Perdikaris, et al. Multiscale modeling meets machine learning: What can we learn? Archives of Computational Methods in Engineering, 28(3):1017–1037, 2021.

[19] George Em Karniadakis, Ioannis G Kevrekidis, Lu Lu, Paris Perdikaris, Sifan Wang, and Liu Yang. Physics-informed machine learning. Nature Reviews Physics, 3(6):422–440, 2021.

[20] Yuntian Chen and Dongxiao Zhang. Integration of knowledge and data in machine learning. arXiv preprint arXiv:2202.10337, 2022.

[21] Alireza Yazdani, Lu Lu, Maziar Raissi, and George Em Karniadakis. Systems biology informed deep learning for inferring parameters and hidden dynamics. PLoS computational biology, 16(11):e1007575, 2020.

[22] George EP Box and HL Lucas. Design of experiments in non-linear situations. Biometrika, 46(1/2):77–90, 1959.

[23] Samuel Bandara, Johannes P. Schlöder, Roland Eils, Hans Georg Bock, and Tobias Meyer. Optimal experimental design for parameter estimation of a cell signaling model. PLOS Computational Biology, 5(11):1–12, 11 2009.

[24] Georg K. Gerber, Andrew B. Onderdonk, and Lynn Bry. Inferring dynamic signatures of microbes in complex host ecosystems. PLOS Computational Biology, 8(8):1–14, 08 2012.

[25] Ali Shahmohammadi and Kimberley B McAuley. Using prior parameter knowledge in modelbased design of experiments for pharmaceutical production. AIChE Journal, 66(11):e17021, 2020.

[26] Juliane Liepe, Sarah Filippi, Michał Komorowski, and Michael P. H. Stumpf. Maximizing the information content of experiments in systems biology. PLOS Computational Biology, 9(1):1–13, 01 2013.

[27] Isabella Verdinelli and Joseph B Kadane. Bayesian designs for maximizing information and outcome. Journal of the American Statistical Association, 87(418):510–515, 1992.

[28] Morris H DeGroot. Uncertainty, information, and sequential experiments. The Annals of Mathematical Statistics, 33(2):404–419, 1962.

[29] Kathryn Chaloner and Isabella Verdinelli. Bayesian experimental design: A review. Statistical Science, 10:273–304, 1995.

[30] Tijana Radivojević, Zak Costello, Kenneth Workman, and Hector Garcia Martin. A machine learning automated recommendation tool for synthetic biology. Nature communications, 11(1):1–14, 2020.

[31] Rémi Lam, Matthias Poloczek, Peter Frazier, and Karen E Willcox. Advances in bayesian optimization with applications in aerospace engineering. In 2018 AIAA Non-Deterministic Approaches Conference, page 1656, 2018.

[32] Daniel James Lizotte. Practical bayesian optimization. University of Alberta, 2008.

[33] Bo Wang and Tao Chen. Gaussian process regression with multiple response variables. Chemometrics and Intelligent Laboratory Systems, 142:159–165, 2015.

[34] George EP Box and Norman R Draper. A basis for the selection of a response surface design. Journal of the American Statistical Association, 54(287):622–654, 1959.

[35] Yael Litvak, Mariana X Byndloss, and Andreas J Bäumler. Colonocyte metabolism shapes the gut microbiota. Science, 362(6418):eaat9076, 2018.

[36] Naschla Gasaly, Marcela A Hermoso, and Martín Gotteland. Butyrate and the fine-tuning of colonic homeostasis: implication for inflammatory bowel diseases. International Journal of Molecular Sciences, 22(6):3061, 2021.

[37] Feng Gao, Yi-Wei Lv, Jie Long, Jie-Mei Chen, Jiu-ming He, Xiong-Zhong Ruan, and Hai-bo Zhu. Butyrate improves the metabolic disorder and gut microbiome dysbiosis in mice induced by a high-fat diet. Frontiers in pharmacology, 10:1040, 2019.

[38] Meire Fortunato, Charles Blundell, and Oriol Vinyals. Bayesian recurrent neural networks. arXiv preprint arXiv:1704.02798, 2017.

[39] Lior Hirschfeld, Kyle Swanson, Kevin Yang, Regina Barzilay, and Connor W Coley. Uncertainty quantification using neural networks for molecular property prediction. Journal of Chemical Information and Modeling, 60(8):3770–3780, 2020.

[40] Yarin Gal and Zoubin Ghahramani. Dropout as a bayesian approximation: Representing model uncertainty in deep learning. In international conference on machine learning, pages 1050–1059. PMLR, 2016.

[41] Armen Der Kiureghian and Ove Ditlevsen. Aleatory or epistemic? does it matter? Structural safety, 31(2):105–112, 2009.

[42] H Bouallagui, Y Touhami, R Ben Cheikh, and M Hamdi. Bioreactor performance in anaerobic digestion of fruit and vegetable wastes. Process biochemistry, 40(3-4):989–995, 2005.

[43] Sherin Varghese, ND Dhanraj, Sharrel Rebello, Raveendran Sindhu, Parameswaran Binod, Ashok Pandey, MS Jisha, and Mukesh Kumar Awasthi. Leads and hurdles to sustainable microbial bioplastic production. Chemosphere, 305:135390, 2022.

[44] Helena Moralejo-Gárate, Emily Mar’Atusalihat, Robbert Kleerebezem, and Mark van Loos-drecht. Microbial community engineering for biopolymer production from glycerol. Applied microbiology and biotechnology, 92(3):631–639, 2011.

[45] Jong Woo Kim, Byung Jun Park, Tae Hoon Oh, and Jong Min Lee. Model-based reinforcement learning and predictive control for two-stage optimal control of fed-batch bioreactor. Computers & Chemical Engineering, 154:107465, 2021.

[46] Kang Zhou, Kangjian Qiao, Steven Edgar, and Gregory Stephanopoulos. Distributing a metabolic pathway among a microbial consortium enhances production of natural products. Nature biotechnology, 33(4):377–383, 2015.

[47] Ryan Tsoi, Feilun Wu, Carolyn Zhang, Sharon Bewick, David Karig, and Lingchong You. Metabolic division of labor in microbial systems. Proceedings of the National Academy of Sciences, 115(10):2526–2531, 2018.

[48] Piotr Oleskowicz-Popiel. Designing reactor microbiomes for chemical production from organic waste. Trends in biotechnology, 36(8):747–750, 2018.

[49] Christopher W Marshall, Edward V LaBelle, and Harold D May. Production of fuels and chemicals from waste by microbiomes. Current opinion in biotechnology, 24(3):391–397, 2013.

[50] Ashley Shade, Hannes Peter, Steven D Allison, Didier L Baho, Mercè Berga, Helmut Bürgmann, David H Huber, Silke Langenheder, Jay T Lennon, Jennifer BH Martiny, et al. Fundamentals of microbial community resistance and resilience. Frontiers in microbiology, 3:417, 2012.

[51] Joshua E Goldford, Nanxi Lu, Djordje Bajić, Sylvie Estrela, Mikhail Tikhonov, Alicia Sanchez-Gorostiaga, Daniel Segrè, Pankaj Mehta, and Alvaro Sanchez. Emergent simplicity in microbial community assembly. Science, 361(6401):469–474, 2018.

[52] Michael L Shuler, Fikret Kargi, and Matthew P DeLisa. Bioprocess Engineering: Basic Concepts. Number 3. Pearson, 2017.

[53] Jean-Loup Faulon and Léon Faure. In silico, in vitro, and in vivo machine learning in synthetic biology and metabolic engineering. Current Opinion in Chemical Biology, 65:85–92, 2021.

[54] Allison J Lopatkin and James J Collins. Predictive biology: modelling, understanding and harnessing microbial complexity. Nature Reviews Microbiology, 18(9):507–520, 2020.

[55] Christopher E Lawson, William R Harcombe, Roland Hatzenpichler, Stephen R Lindemann, Frank E Löffler, Michelle A O’Malley, Héctor García Martín, Brian F Pfleger, Lutgarde Raskin, Ophelia S Venturelli, et al. Common principles and best practices for engineering microbiomes. Nature Reviews Microbiology, 17(12):725–741, 2019.

[56] Karsten Zengler, Kirsten Hofmockel, Nitin S Baliga, Scott W Behie, Hans C Bernstein, James B Brown, José R Dinneny, Sheri A Floge, Samuel P Forry, Matthias Hess, et al. Ecofabs: advancing microbiome science through standardized fabricated ecosystems. Nature methods, 16(7):567–571, 2019.

[57] David A Cohn, Zoubin Ghahramani, and Michael I Jordan. Active learning with statistical models. Journal of artificial intelligence research, 4:129–145, 1996.

[58] David JC MacKay. Information-based objective functions for active data selection. Neural computation, 4(4):590–604, 1992.

[59] Marco Tulio Ribeiro, Sameer Singh, and Carlos Guestrin. “why should i trust you?” explaining the predictions of any classifier. In Proceedings of the 22nd ACM SIGKDD international conference on knowledge discovery and data mining, pages 1135–1144, 2016.

[60] Ricky TQ Chen, Yulia Rubanova, Jesse Bettencourt, and David K Duvenaud. Neural ordinary differential equations. Advances in neural information processing systems, 31, 2018.

[61] Michael E Tipping. Sparse bayesian learning and the relevance vector machine. Journal of machine learning research, 1(Jun):211–244, 2001.

[62] Brian Munsky, William S Hlavacek, and Lev S Tsimring. Quantitative biology: theory, computational methods, and models. MIT Press, 2018.

[63] SP Luttrell. The use of transinformation in the design of data sampling schemes for inverse problems. Inverse Problems, 1(3):199, 1985.

[64] Kapil G Gadkar, Rudiyanto Gunawan, and Francis J Doyle. Iterative approach to model identification of biological networks. BMC bioinformatics, 6(1):1–20, 2005.

[65] Gaia Franceschini and Sandro Macchietto. Model-based design of experiments for parameter precision: State of the art. Chemical Engineering Science, 63(19):4846–4872, 2008.

