## Supplementary Information for "Integrating a tailored recurrent neural network with Bayesian experimental design to optimize microbial community functions"

Jaron C. Thompson[1,2], Victor M. Zavala[1], and Ophelia S. Venturelli[1,2,3*]
[1]Department of Chemical and Biological Engineering
[2]Department of Biochemistry
[3]Department of Bacteriology
University of Wisconsin-Madison, Madison, Wisconsin, 53706

#### 1 SI Text

##### 1.1 Data pre-processing

To train the MiRNN, the data are pre-processed such that each species and each metabolite are first normalized by their maximum value in the training data. Given un-normalized training data, $\hat{\mathcal{D}}(\mathbf{q}) = \{\hat{\mathbf{y}}(q_1), ..., \hat{\mathbf{y}}(q_n)\}$, each output, indexed by $j = 1, ..., n_y$, is normalized so that

$$y_j(q_i) = \frac{\hat{y}_j(q_i)}{\max\{\hat{y}_j(q_k)\}_{k=1}^{n}} = \frac{\hat{y}_j(q_i)}{c_j} \tag{1}$$

where $\hat{y}_j(q_i)$ is the un-normalized (measured) output $j$ from experimental condition $q_i$, $y_j(q_i)$ is the normalized output variable $j$ from experimental condition $q_i$, and $c_j$ is the maximum value of species $j$ found in $\hat{\mathcal{D}}(\mathbf{q})$. When making predictions for any experimental condition $q_i$, the inverse transform is applied to the model output so that

$$\mathbb{E}[\hat{\mathbf{y}}(q_i)] = \text{diag}(\mathbf{c}) \cdot \mathcal{M}(\theta, q_i) \tag{2}$$

$$\text{Var}[\hat{\mathbf{y}}(q_i)] = \text{diag}(\mathbf{c}) \cdot \mathbf{\Sigma}_y(q_i) \cdot \text{diag}(\mathbf{c}) \tag{3}$$

##### 1.2 Evaluation of model prediction performance

###### 1.2.1 Pearson correlation coefficient

The Pearson correlation coefficient ($R$) was computed using the LINREGRESS function from SCIPY [9]. The Pearson correlation coefficient corresponding to prediction performance of output $j$ over an un-normalized test data set, $\hat{\mathcal{D}}(\mathbf{q}) = \{\hat{\mathbf{y}}(q_1), ..., \hat{\mathbf{y}}(q_n)\}$ is given by

$$R_j = \frac{\sum_{i=1}^{n}(\hat{y}_j(q_i) - \frac{1}{n}\sum_{k=1}^{n}\hat{y}_j(q_k))(\mathbb{E}[\hat{y}_j(q_i)] - \frac{1}{n}\sum_{k=1}^{n}\mathbb{E}[\hat{y}_j(q_k)])}{\sqrt{\sum_{i=1}^{n}(\hat{y}_j(q_i) - \frac{1}{n}\sum_{k=1}^{n}\hat{y}_j(q_k))^2 \sum_{i=1}^{n}(\mathbb{E}[\hat{y}_j(q_i)] - \frac{1}{n}\sum_{k=1}^{n}\mathbb{E}[\hat{y}_j(q_k)])^2}} \tag{4}$$

###### 1.2.2 Mean squared error

The mean square error between measured values and model predictions of output $j$ is given by

$$MSE_j = \frac{1}{n}\sum_{i=1}^{n}(\hat{y}_j(q_i) - \mathbb{E}[\hat{y}_j(q_i)])^2. \tag{5}$$

###### 1.2.3 Log-likelihood of test data

The log-likelihood is the log of the probability of a data set given the model. The log-likelihood evaluated for a data set, $\mathcal{D}(\mathbf{q})$, using model predicted covariance is given by

$$\begin{aligned}\ln p(\mathcal{D}(\mathbf{q})|\mathcal{M}(\theta,\mathbf{q}), \mathbf{\Sigma}_y(\mathbf{q})) = &-\sum_{i=1}^{n}(\mathbf{y}(q_i) - \mathcal{M}(\theta, q_i))^T \cdot \mathbf{\Sigma}_{\mathbf{y}}(q_i) \cdot (\mathbf{y}(q_i) - \mathcal{M}(\theta, q_i)) \\ &-\sum_{i=1}^{n}\frac{1}{2}\ln \det \mathbf{\Sigma}_{\mathbf{y}}(q_i) - \frac{n \cdot n_y}{2}\ln(2\pi)\end{aligned} \tag{6}$$

The log-likelihood evaluated using fixed covariance is given by

$$\begin{aligned}\ln p(\mathcal{D}(\mathbf{q})|\mathcal{M}(\theta,\mathbf{q}), \mathbf{\Sigma}_y) = &-\sum_{i=1}^{n}(\mathbf{y}(q_i) - \mathcal{M}(\theta, q_i))^T \cdot \mathbf{\Sigma}_{\mathbf{y}} \cdot (\mathbf{y}(q_i) - \mathcal{M}(\theta, q_i)) \\ &-\sum_{i=1}^{n}\frac{1}{2}\ln \det \mathbf{\Sigma}_{\mathbf{y}} - \frac{n \cdot n_y}{2}\ln(2\pi)\end{aligned} \tag{7}$$

##### 1.3 Hyper-parameter optimization

Optimization of hyper-parameters using only the training data is often called *empirical Bayes* [1]. We use an empirical Bayes framework where model parameters are optimized using a Gauss-Newton algorithm and the parameter posterior density is approximated using the Laplace approximation [3]. To determine a set of optimal hyper-parameters, $\xi^* = \{\mathbf{\Sigma}_\theta^*, \mathbf{\Sigma}_y^*\}$, we seek $\xi$ that maximizes the marginal likelihood function given by

$$p(\mathcal{D}(\mathbf{q})|\xi) = \int_\theta p(\mathcal{D}(\mathbf{q}), \theta|\xi)d\theta. \tag{8}$$

We can use the *expectation maximization* (EM) algorithm to update $\xi^{(l+1)}$, which involves maximizing the expected log likelihood,

$$\xi^{(l+1)} = \underset{\xi}{\text{argmax}}\ \mathbb{E}_{\theta|\mathcal{D}(\mathbf{q}),\xi^{(l)}}[\ln p(\mathcal{D}(\mathbf{q}), \theta|\xi)], \tag{9}$$

followed by re-evaluation of the posterior parameter distribution $p(\theta|\mathcal{D}(\mathbf{q}), \xi^{(l+1)})$. The process of maximizing Eq 9 and updating the posterior parameter distribution is repeated until convergence of the marginal likelihood function given by Eq 14.

Given an initial guess for the hyper-parameters, $\xi^{(l)}$, the posterior parameter distribution is approximated following the steps described in the methods section, *Bayesian estimation and uncertainty quantification*. With $\ln p(\mathcal{D}(\mathbf{q}), \theta) = \ln p(\mathcal{D}(\mathbf{q})|\theta) + \ln p(\theta)$, Eq 9 becomes

$$\mathbb{E}_{\theta|\mathcal{D}(\mathbf{q}),\xi^{(l)}}[\ln p(\mathcal{D}(\mathbf{q}), \theta|\xi)] = \mathbb{E}_{\theta|\mathcal{D}(\mathbf{q}),\xi^{(l)}}[\ln p(\mathcal{D}(\mathbf{q})|\theta, \xi)] + \mathbb{E}_{\theta|\mathcal{D}(\mathbf{q}),\xi^{(l)}}[\ln p(\theta|\xi)]. \tag{10}$$

The first term is the expectation of the log likelihood,

$$\begin{aligned}\mathbb{E}_{\theta|\mathcal{D}(\mathbf{q}),\xi^{(l)}}[\ln p(\mathcal{D}(\mathbf{q})|\theta, \xi)] = &-\frac{1}{2}\sum_{i=1}^{n}\mathbb{E}_{\theta|\mathcal{D}(\mathbf{q}),\xi^{(l)}}\left[(\mathbf{y}(q_i) - \mathcal{M}(\theta, q_i))^T \mathbf{\Sigma}_y^{-1}(\mathbf{y}(q_i) - \mathcal{M}(\theta, q_i))\right] \\ &- \frac{|\mathcal{D}(\mathbf{q})|}{2}\ln\left(2\pi \det \mathbf{\Sigma}_y\right).\end{aligned}$$

linearizing the model with respect to $\theta$ about $\theta_{\mathrm{MAP}}$ and evaluating the expectation gives

$$\begin{aligned}\mathbb{E}_{\theta|\mathcal{D}(\mathbf{q}),\xi^{(l)}}[\ln p(\mathcal{D}(\mathbf{q})|\theta, \xi)] = &-\frac{1}{2}\sum_{i=1}^{n}(\mathbf{y}(q_i) - \mathcal{M}(\theta_{\mathrm{MAP}}, q_i))^T\, \mathbf{\Sigma}_y^{-1}\,(\mathbf{y}(q_i) - \mathcal{M}(\theta_{\mathrm{MAP}}, q_i)) \\ &+ \mathrm{Tr}\left(\mathbf{\Sigma}_y^{-1}\mathbf{G}(\mathbf{q}, q_i)\mathbf{H}(\mathbf{q})^{-1}\mathbf{G}(\mathbf{q}, q_i)^T\right) - \frac{|\mathcal{D}(\mathbf{q})|}{2}\ln\left(2\pi \det \mathbf{\Sigma}_y\right).\end{aligned}$$

Evaluating the second term in Eq 10 gives

$$\mathbb{E}_{\theta|\mathcal{D}(\mathbf{q}),\xi^{(l)}}[\ln p(\theta|\xi)] = -\frac{1}{2}\sum_{k=1}^{n_\theta}\left(\ln\,[\alpha]_k - [\alpha]_k \cdot ([\theta_{\mathrm{MAP}}^2]_k + [\mathbf{H}(\mathbf{q})^{-1}]_{kk}) - \ln\,2\pi\right)$$

Update equations for $\xi$ are found by taking the derivative of Eq 10 with respect to $\mathbf{\Sigma}_y$ and $[\alpha]_k$ and solving for $\mathbf{\Sigma}_y^{(l+1)}$ and $[\alpha^{(l+1)}]_k$,

$$\begin{aligned}\mathbf{\Sigma}_y^{(l+1)} = &\frac{1}{|\mathcal{D}(\mathbf{q})|}\sum_{i=1}^{n}(\mathbf{y}(q_i) - \mathcal{M}(\theta_{\mathrm{MAP}}, q_i))^T \mathbf{\Sigma}_y^{(l)\;-1}(\mathbf{y}(q_i) - \mathcal{M}(\theta_{\mathrm{MAP}}, q_i)) \\ &+ \mathbf{G}(\mathbf{q}, q_i)\mathbf{H}(\mathbf{q})^{-1}\mathbf{G}(\mathbf{q}, q_i)^T,\end{aligned} \tag{11}$$

$$[\alpha^{(l+1)}]_k = \frac{1}{\left([\theta_{\mathrm{MAP}}^2]_k + [\mathbf{H}(\mathbf{q})^{-1}]_{kk}\right)}. \tag{12}$$

Model hyper-parameters are updated until convergence of the log of the model evidence (i.e. marginal likelihood), which is approximated using the Laplace approximation as

$$\ln\,p(\mathcal{D}(\mathbf{q})|\xi) \approx -\frac{1}{2}\ln\,\det\,\mathbf{\Sigma}_\theta - \frac{|\mathcal{D}(\mathbf{q})|}{2}\ln\,2\pi\,\det\,\mathbf{\Sigma}_y - \frac{1}{2}\ln\,\det\,\mathbf{H}(\mathbf{q}) - \theta_{\mathrm{MAP}}^T\mathbf{\Sigma}_y^{-1}\theta_{\mathrm{MAP}} \tag{13}$$

$$-\frac{1}{2}\sum_{i=1}^{n}(\mathbf{y}(q_i) - \mathcal{M}(\theta_{\mathrm{MAP}}, q_i))^T\mathbf{\Sigma}_y^{-1}(\mathbf{y}(q_i) - \mathcal{M}(\theta_{\mathrm{MAP}}, q_i)). \tag{14}$$

#### 1.4 Function to quantify information content

Given a model that has been trained on previous data, $\mathcal{D}(\mathbf{q}^{(l)})$, where $\mathbf{q}^{(l)}$ represents all previous experimental designs, we wish to design the next experiment, $\mathbf{q}^{(l+1)}$. Using principles of Bayesian experimental design, the information content of the next experiment $\mathbf{q}^{(l+1)} \subset Q$ is evaluated using the expected gain in information that would result from updating the current model with a new dataset $\mathcal{D}(\mathbf{q}^{(l+1)}) = \{\mathbf{y}(q_1^{(l+1)}), ..., \mathbf{y}(q_n^{(l+1)})\}$ [8, 6]. The expected gain in information is the expected Kullback-Leibler divergence between the parameter posterior and current distribution, which is denoted as $p(\theta)$ in place of $p(\theta|\mathcal{D}(\mathbf{q}^{(l)}))$ to simplify the notation.

$$
\begin{aligned}
f_I(\mathbf{q}^{(l)}, \mathbf{q}^{(l+l)}) &:= \mathbb{E}_{\mathcal{D}(\mathbf{q}^{(l+l)})}[\mathrm{KL}\left(p(\theta|\mathcal{D}(\mathbf{q}^{(l+l)}))||p(\theta)\right)] \\
&= \int_{\mathcal{D}(\mathbf{q}^{(l+l)})} \int_\theta p(\theta|\mathcal{D}(\mathbf{q}^{(l+l)})) \ln\left(\frac{p(\theta|\mathcal{D}(\mathbf{q}^{(l+l)}))}{p(\theta)}\right) \mathrm{d}\theta \; p(\mathcal{D}(\mathbf{q}^{(l+l)})) \; \mathrm{d}\mathcal{D}(\mathbf{q}^{(l+l)}) \\
&= \int_{\mathcal{D}(\mathbf{q}^{(l+l)})} \int_\theta p(\theta|\mathcal{D}(\mathbf{q}^{(l+l)})) \ln\left(p(\theta|\mathcal{D}(\mathbf{q}^{(l+l)}))\right) \mathrm{d}\theta \; p(\mathcal{D}(\mathbf{q}^{(l+l)})) \; \mathrm{d}\mathcal{D}(\mathbf{q}^{(l+l)}) \\
&\quad - \int_{\mathcal{D}(\mathbf{q}^{(l+l)})} \int_\theta p(\theta|\mathcal{D}(\mathbf{q}^{(l+l)})) \ln\left(p(\theta)\right) \mathrm{d}\theta \; p(\mathcal{D}(\mathbf{q}^{(l+l)})) \; \mathrm{d}\mathcal{D}(\mathbf{q}^{(l+l)}) \\
&= \int_{\mathcal{D}(\mathbf{q}^{(l+l)})} \int_\theta \ln\left(p(\theta|\mathcal{D}(\mathbf{q}^{(l+l)}))\right) p(\theta|\mathcal{D}(\mathbf{q}^{(l+l)})) p(\mathcal{D}(\mathbf{q}^{(l+l)})) \mathrm{d}\theta \; \mathrm{d}\mathcal{D}(\mathbf{q}^{(l+l)}) \\
&\quad - \int_\theta \ln\left(p(\theta)\right) \int_{\mathcal{D}(\mathbf{q}^{(l+l)})} p(\theta|\mathcal{D}(\mathbf{q}^{(l+l)})) p(\mathcal{D}(\mathbf{q}^{(l+l)})) \mathrm{d}\mathcal{D}(\mathbf{q}^{(l+l)}) \mathrm{d}\theta \\
&= \int_{\mathcal{D}(\mathbf{q}^{(l+l)})} \int_\theta \ln\left(p(\theta|\mathcal{D}(\mathbf{q}^{(l+l)}))\right) p(\theta, \mathcal{D}(\mathbf{q}^{(l+l)})) \mathrm{d}\theta \; \mathrm{d}\mathcal{D}(\mathbf{q}^{(l+l)}) - \int_\theta \ln\left(p(\theta)\right) p(\theta) \mathrm{d}\theta \\
&= -h\left[\theta|\mathcal{D}(\mathbf{q}^{(l+l)})\right] + h\left[\theta\right]
\end{aligned} \tag{15}
$$

where $h\left[\theta\right]$ is the entropy of the current parameter distribution and $h\left[\theta|\mathcal{D}(\mathbf{q}^{(l+l)})\right]$ is the conditional entropy of the parameter distribution given the dataset $\mathcal{D}(\mathbf{q}^{(l+l)})$. Interpreting entropy as a measure of uncertainty, the expected gain in information quantifies the amount the model expects the data, $\mathcal{D}(\mathbf{q}^{(l+l)})$, to reduce uncertainty in the model's current parameter values. Assuming that the posterior parameter distribution conditioned on $\mathcal{D}(\mathbf{q}^{(l+l)})$ is Gaussian with precision matrix, $\mathbf{H}(\mathbf{q}^{(l)}, \mathbf{q}^{(l+l)})$, we can analytically evaluate the integral over $\theta$. Keeping only terms that depend on $\mathbf{q}^{(l)}$ and $\mathbf{q}^{(l+l)}$, this gives

$$
\begin{aligned}
-h[\theta|\mathcal{D}(\mathbf{q}^{(l+l)}))] &= \int_{\mathcal{D}(\mathbf{q}^{(l+l)})} \int_\theta \ln\left(p(\theta|\mathcal{D}(\mathbf{q}^{(l+l)}))\right) p(\theta|\mathcal{D}(\mathbf{q}^{(l+l)})) \mathrm{d}\theta \; p(\mathcal{D}(\mathbf{q}^{(l+l)})) \; \mathrm{d}\mathcal{D}(\mathbf{q}^{(l+l)}) \\
&= \int_{\mathcal{D}(\mathbf{q}^{(l+l)})} \ln\det \mathbf{H}(\mathbf{q}^{(l)}, \mathbf{q}^{(l+l)}) p(\mathcal{D}(\mathbf{q}^{(l+l)})) \; \mathrm{d}\mathcal{D}(\mathbf{q}^{(l+l)})
\end{aligned} \tag{16}
$$

We can once again make use of the Laplace approximation to get an expression for the posterior precision matrix, $\mathbf{H}(\mathbf{q}^{(l)}, \mathbf{q}^{(l+l)})$. Starting with Bayes' theorem, the parameter distribution conditioned on $\mathcal{D}(\mathbf{q}^{(l+l)})$ is proportional to the product of the likelihood of the data multiplied by the current parameter distribution

$$p(\theta|\mathcal{D}(\mathbf{q}^{(l+l)})) \propto p(\mathcal{D}(\mathbf{q}^{(l+l)})|\theta)p(\theta),$$

where $p(\theta) = \mathcal{N}(\theta_{\mathrm{MAP}}(\mathbf{q}), \mathbf{H}(\mathbf{q})^{-1})$. The Laplace approximation of the posterior parameter precision matrix is determined by taking the Hessian of the negative log of this posterior parameter distribution,

$$\begin{aligned}
\mathbf{H}(\mathbf{q}^{(l)}, \mathbf{q}^{(l+l)}) &= -\nabla_\theta \nabla_\theta \ln p(\theta) - \nabla_\theta \nabla_\theta \ln p(\mathcal{D}(\mathbf{q}^{(l+l)})|\theta) \\
&= \mathbf{H}(\mathbf{q}^{(l)}) + \frac{1}{2}\nabla_\theta \nabla_\theta \sum_{i=1}^{n} (\mathbf{y}(q_i) - \mathcal{M}(\theta, q_i^{(l+l)}))^T \mathbf{\Sigma}_y^{-1} (\mathbf{y}(q_i) - \mathcal{M}(\theta, q_i^{(l+l)})) \\
&= \mathbf{H}(\mathbf{q}^{(l)}) + \sum_{i=1}^{n} \mathbf{G}(\mathbf{q}^{(l)}, q_i^{(l+l)})^T \mathbf{\Sigma}_y^{-1} \mathbf{G}(\mathbf{q}^{(l)}, q_i^{(l+l)}) \\
&\qquad + (\mathcal{M}(\theta, q_i^{(l+l)}) - \mathbf{y}(q_i))^T \mathbf{\Sigma}_y^{-1} \nabla_\theta \nabla_\theta \mathcal{M}(\theta, q_i^{(l+l)}).
\end{aligned}$$

Because $\mathbf{y}(q_i)$ is modeled as a Gaussian with mean given by the model prediction, $\mathcal{M}(\theta, q_i^{(l+l)})$, for each $\mathbf{y}(q_i) \in \mathcal{D}(\mathbf{q}^{(l+l)})$, the summation over the residuals, $\mathcal{M}(\theta, q_i^{(l+l)}) - \mathbf{y}(q_i)$, vanishes when evaluating the expectation over $\mathcal{D}(\mathbf{q}^{(l+l)})$ in Eq. 16, assuming that the residuals are uncorrelated with the second derivative of the model with respect to parameters[1]. Evaluating Eq. 16 gives the final expression for the information function

$$\mathrm{EIG}(\mathbf{q}^{(l)}, \mathbf{q}^{(l+l)}) \approx \ln\det\left(\mathbf{H}(\mathbf{q}^{(l)}) + \sum_{i=1}^{n} \mathbf{G}(\mathbf{q}^{(l)}, q_i^{(l+l)})^T \mathbf{\Sigma}_y^{-1} \mathbf{G}(\mathbf{q}^{(l)}, q_i^{(l+l)})\right) - \ln\det\left(\mathbf{H}(\mathbf{q}^{(l)})\right). \quad (17)$$

Experimental designs that maximize Eq. 17 are called Bayesian D-optimal[7, 2].

##### 1.5 Fast evaluation of the information function

Evaluation of the function given by Eq. 17 can be computationally expensive for models with a large number of parameters. Alternatively, we can compute the EIG using an equivalent expression,

$$\begin{aligned}
\mathrm{EIG}(\mathbf{q}^{(l)}, \mathbf{q}^{(l+l)}) &\approx \ln\det\left(\mathbf{H}(\mathbf{q}^{(l)}) + \sum_{i=1}^{n} \mathbf{G}(\mathbf{q}^{(l)}, q_i^{(l+l)})^T \mathbf{\Sigma}_y^{-1} \mathbf{G}(\mathbf{q}^{(l)}, q_i^{(l+l)})\right) - \ln\det\left(\mathbf{H}(\mathbf{q}^{(l)})\right) \\
&= \sum_{i=1}^{n} \ln\det\left(\mathbb{I}_{n_y} + \mathbf{\Sigma}_y^{-1} \mathbf{G}(\mathbf{q}^{(l)}, q_i^{(l+l)}) \mathbf{A}_{i-1}^{-1} \mathbf{G}(\mathbf{q}^{(l)}, q_i^{(l+l)})^T\right) \quad (18)
\end{aligned}$$

where

$$\mathbf{A}_i = \mathbf{A}_{i-1} + \mathbf{G}(\mathbf{q}^{(l)}, q_i^{(l+l)})^T \mathbf{\Sigma}_y^{-1} \mathbf{G}(\mathbf{q}^{(l)}, q_i^{(l+l)}), \qquad \mathbf{A}_0 = \mathbf{H}(\mathbf{q}^{(l)}). \quad (19)$$

Consequently, we can avoid taking the determinant of a matrix whose dimension is $n_\theta \times n_\theta$ in favor of evaluating the determinant and inverse of $n$ matrices each with dimension $n_y \times n_y$ given $\mathbf{H}(\mathbf{q}^{(l)})^{-1}$. The matrix inverse $\mathbf{A}_i^{-1}$ can be evaluated efficiently using the Woodbury identity,

$$\mathbf{A}_i^{-1} = \mathbf{A}_{i-1}^{-1} - \mathbf{A}_{i-1}^{-1}\mathbf{G}(\mathbf{q}^{(l)}, q_i^{(l+1)})^T(\boldsymbol{\Sigma}_y + \mathbf{G}(\mathbf{q}^{(l)}, q_i^{(l+1)})\mathbf{A}_{i-1}^{-1}\mathbf{G}(\mathbf{q}^{(l)}, q_i^{(l+1)})^T)^{-1}\mathbf{G}(\mathbf{q}^{(l)}, q_i^{(l+1)})\mathbf{A}_{i-1}^{-1} \tag{20}$$

To see that the two expressions for the EIG in Eq. 18 are equivalent, we start with the following identity [1],

$$\det(\mathbb{I}_{n_\theta} + \mathbf{X}\mathbf{Z}^T) = \det(\mathbb{I}_{n_y} + \mathbf{X}^T\mathbf{Z}) \tag{21}$$

where $\mathbf{X}$ and $\mathbf{Z}$ have dimensions $n_\theta \times n_y$. Replacing terms with $\mathbf{X} = \mathbf{A}_{i-1}^{-1}\mathbf{G}(\mathbf{q}^{(l)}, q_i^{(l+1)})^T\boldsymbol{\Sigma}_y^{-1}$ and $\mathbf{Z}^T = \mathbf{G}(\mathbf{q}^{(l)}, q_i^{(l+1)})$, we get

$$\det(\mathbb{I}_{n_\theta} + \mathbf{A}_{i-1}^{-1}\mathbf{G}(\mathbf{q}^{(l)}, q_i^{(l+1)})^T\boldsymbol{\Sigma}_y^{-1}\mathbf{G}(\mathbf{q}^{(l)}, q_i^{(l+1)})) = \det(\mathbb{I}_{n_y} + \boldsymbol{\Sigma}_y^{-1}\mathbf{G}(\mathbf{q}^{(l)}, q_i^{(l+1)})\mathbf{A}_{i-1}^{-1}\mathbf{G}(\mathbf{q}^{(l)}, q_i^{(l+1)})^T). \tag{22}$$

Using $\det(\mathbf{XZ}) = \det(\mathbf{X})\det(\mathbf{Z})$, and multiplying both sides of Eq. 22 by $\det(\mathbf{A}_{i-1})$, we have

$$\begin{aligned}&\det(\mathbf{A}_{i-1} + \mathbf{G}(\mathbf{q}^{(l)}, q_i^{(l+1)})^T\boldsymbol{\Sigma}_y^{-1}\mathbf{G}(\mathbf{q}^{(l)}, q_i^{(l+1)}))\\ &\qquad = \det(\mathbf{A}_{i-1})\det(\mathbb{I}_{n_y} + \boldsymbol{\Sigma}_y^{-1}\mathbf{G}(\mathbf{q}^{(l)}, q_i^{(l+1)})\mathbf{A}_{i-1}^{-1}\mathbf{G}(\mathbf{q}^{(l)}, q_i^{(l+1)})^T)\end{aligned} \tag{23}$$

Using $\mathbf{A}_i = \mathbf{A}_{i-1} + \mathbf{G}(\mathbf{q}^{(l)}, q_i^{(l+l)})^T\boldsymbol{\Sigma}_y^{-1}\mathbf{G}(\mathbf{q}^{(l)}, q_i^{(l+l)})$, we can express the summation in Eq. 18 as

$$\mathbf{H}(\mathbf{q}^{(l)}) + \sum_{i=1}^{n}\mathbf{G}(\mathbf{q}^{(l)}, q_i^{(l+l)})^T\boldsymbol{\Sigma}_y^{-1}\mathbf{G}(\mathbf{q}^{(l)}, q_i^{(l+l)}) = \mathbf{A}_{n-1} + \mathbf{G}(\mathbf{q}^{(l)}, q_n^{(l+l)})^T\boldsymbol{\Sigma}_y^{-1}\mathbf{G}(\mathbf{q}^{(l)}, q_n^{(l+l)}). \tag{24}$$

Using the identity in Eq. 23, we have

$$\begin{aligned}&\det(\mathbf{A}_{n-1} + \mathbf{G}(\mathbf{q}^{(l)}, q_n^{(l+1)})^T\boldsymbol{\Sigma}_y^{-1}\mathbf{G}(\mathbf{q}^{(l)}, q_n^{(l+1)}))\\ &\qquad = \det(\mathbf{A}_{n-1})\det(\mathbb{I}_{n_y} + \boldsymbol{\Sigma}_y^{-1}\mathbf{G}(\mathbf{q}^{(l)}, q_n^{(l+1)})\mathbf{A}_{n-1}^{-1}\mathbf{G}(\mathbf{q}^{(l)}, q_n^{(l+1)})^T).\end{aligned} \tag{25}$$

We can continue to apply Eq. 23 to $\mathbf{A}_{n-1} = \mathbf{A}_{n-2} + \mathbf{G}(\mathbf{q}^{(l)}, q_{n-1}^{(l+l)})^T\boldsymbol{\Sigma}_y^{-1}\mathbf{G}(\mathbf{q}^{(l)}, q_{n-1}^{(l+l)})$ until $n = 0$ and take the natural log of the result to yield Eq. 18.

#### 1.6 Specification of ground truth bioreactor model parameters

The governing equations of the model are given by

$$\begin{aligned}\frac{\mathrm{d}V}{\mathrm{d}t} &= u\\ \frac{\mathrm{d}\mathbf{r}}{\mathrm{d}t} &= \mathbf{r}\odot\left(-\mathbf{s}^T\cdot\mathbf{C} - \mathbf{d}\right) + \frac{u}{V}(\mathbf{r}_f - \mathbf{r})\\ \frac{\mathrm{d}\mathbf{s}}{\mathrm{d}t} &= \mathbf{s}\odot(\mathbf{C}\cdot\mathbf{r} - \mathbf{g}) - \frac{u}{V}\cdot\mathbf{s}\\ \frac{\mathrm{d}m}{\mathrm{d}t} &= \mathbf{y}_{m/s}^T\cdot\frac{\mathrm{d}\mathbf{s}}{\mathrm{d}t_+} - k_d\, m - \frac{u}{V}\, m\end{aligned}$$

where $\mathbf{r}$ is a vector of resource concentrations in the reactor, $\mathbf{r}_f$ is a vector of resource concentrations in the feed, $\mathbf{s}$ is a vector of consumer species, $\mathbf{d}$ is a vector of resource degradation rates, $\mathbf{g}$ is a vector of minimum growth rates needed for each species to survive, $m$ is the metabolite concentration, $\mathbf{y}_{m/s}$ is a vector of yield coefficients, $k_d$ is the product degradation rate, $[\mathbf{C}]_{ij}$ is the rate species $i$ consumes resource $j$, and $u(t)$ represents the rate at which the feed is added to the reactor.

The parameters of the model that need to be specified include $\mathbf{C}$, $\mathbf{d}$, $\mathbf{g}$, $\mathbf{y}_{m/s}$, and $k_d$. The specification of the consumer resource component of the model and its parameters is a modified version of the model presented in [5]. The matrix of species-resource interaction coefficients was determined by first specifying the probability that a species depends on a resource, $p_{s/r}$, which was set to .6 for the simulation. A matrix of *concentration parameters*, denoted as $\mathbf{\Theta}$ determines the degree that a species depends on the concentration of a resource, where

$$[\mathbf{\Theta}]_{ij} = [\mathbf{\Theta}']_{ij} / \sum_{j=1}^{n_r} [\mathbf{\Theta}']_{ij}$$

where $[\mathbf{\Theta}']_{i,j} \sim \text{Uniform}(0,1)$ with probability $p_{s/r}$ and is zero otherwise. The matrix of interaction coefficients are sampled from a Normal distribution with parameters given by

$$[\mathbf{C}]_{ij} \sim \mathcal{N}(\mu = [\mathbf{\Theta}]_{ij}, \sigma = [\mathbf{\Theta}]_{ij}/10).$$

All elements of the degradation rate of metabolites, $\mathbf{d}$, and the minimum amount of resources necessary, $\mathbf{g}$, were set to .01. The degradation rate of the product, $k_d$, was set to .005. The metabolite yield coefficients were specified to be $\mathbf{y}_{m/s} = [0, .5, 0, 0, 0]$, since species two depended on the fewest number of resources. This specification is important so that the optimal set of resources in the feed is not simply the inclusion of all resources.

###### 1.6.1 Specification of experimental design space

We build on the dynamic design of experiments approach outlined in [4] to propose candidate feed-flowrate profiles as a linear combination of orthogonal basis functions. We note that while this is a useful approach for generating feasible control profiles, it is not a requirement for the presented method since the MiRNN is directly a function of the control profile and not the parameters that encode the profile. The batch time for the reactor was set to 130 hours, the initial reactor volume is set to 7L, and a limit on the reactor volume is set to 10L. The limit on reactor volume imposes a constraint on the possible feed-flowrate profiles where

$$V(0) + \int_0^{t_{\text{batch}}} u(t)\text{d}t \leq V_{\max}. \tag{26}$$

By defining a dimensionless time, $\tau = t/t_{\text{batch}}$, and following the steps outlined in [4], we use the following equation to encode different time dependent feed-flowrates,

$$u(\tau) = \frac{6}{130}(1-\tau)(1 + x_1 P_0(\tau) + x_2 P_1(\tau) - (x_1 + x_2) P_2(\tau)), \tag{27}$$

where $P_i(\tau)$ is the $i^{th}$ Legendre polynomial and $-.5 \leq x_1 \pm x_2 \leq .5$. Selecting a different set of the coefficients, $x_i$, results in different feasible feed-flowrate profiles. A set of 20 different feed flow rates was generated using latin-hypercube sampling of each $x_i$. The design space, $Q$, was composed of matching each of the 20 feed flow rates with every possible combination of resources (excluding no resources), which resulted in a set of $20 \times (2^7 - 1) = 2,540$ experimental conditions. Each output in each simulated condition was corrupted with 5% Gaussian noise to mimic variation in experimental measurements.

#### 2 SI Figures

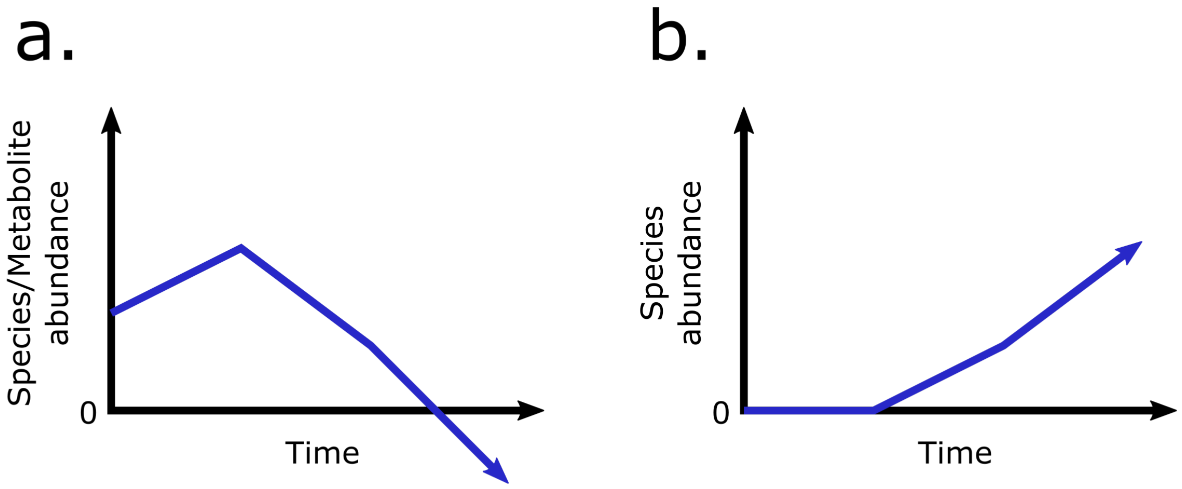


Figure 1: **Examples of physically unrealistic species and metabolite predictions.** (a.) Species and metabolite abundance cannot be negative. (b.) If a species is initially at zero abundance (i.e. not present), it cannot have a positive abundance at later time points.

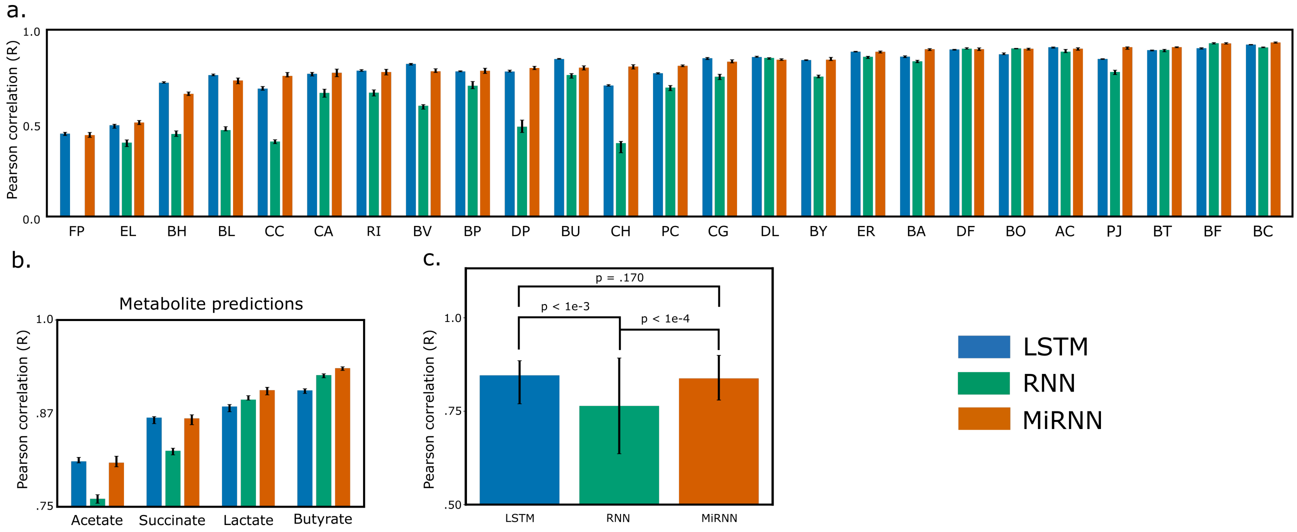


Figure 2: **Comparison of K-fold cross-validation performance.** (a.) Comparison of LSTM, RNN, and MiRNN prediction performance (coefficient of determination) of species abundances after performing 20-fold cross-validation over 10 trials, with the order of samples shuffled in each trial. Bar plot heights indicate the median prediction performance and error bars indicate the interquartile range computed over the 10 trials. (b.) Comparison of LSTM, RNN, and MiRNN prediction performance (coefficient of determination) of metabolite concentrations after performing 20-fold cross-validation over 10 trials, with the order of samples shuffled in each trial. Bar plot heights indicate the median prediction performance and error bars indicate the interquartile range computed over the 10 trials. (c.) A two-tailed paired t-test was used to test for significant differences in the median prediction performance of each species and each metabolite (n=29). The LSTM significantly outperformed the RNN ($p < 1 \times 10^{-3}$) and was slightly outperformed by the MiRNN (p = .170).

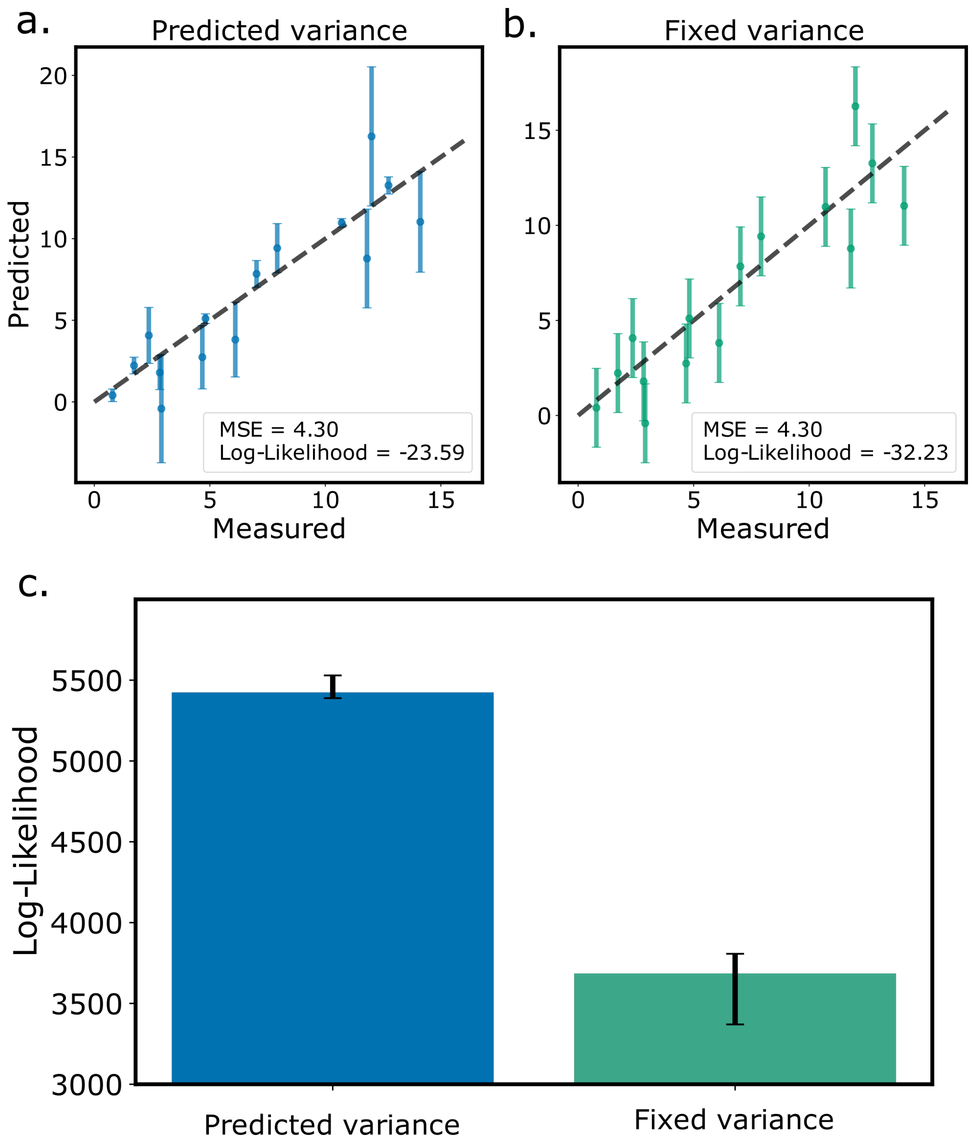


Figure 3: **Illustration of test data log-likelihood.** Comparison of test data log-likelihood using predicted variance versus fixed variance. When deviations between measured and predicted values are high, a corresponding high prediction variance will improve the log-likelihood. Conversely, if deviations between measured and predicted values are small, then a small variance will improve the log-likelihood. (a.) The predicted variance captures variation between measured and predicted values resulting in a higher log-likelihood compared to panel (b.) where prediction uncertainty is based on a fixed estimate of the variance. (c.) Comparison of test data log-likelihood using predicted covariance (left) and fixed covariance (right) after performing 20-fold cross-validation over 10 trials. Bar plot heights indicate the median test data log-likelihood and error bars indicate the interquartile range computed over the 10 trials.

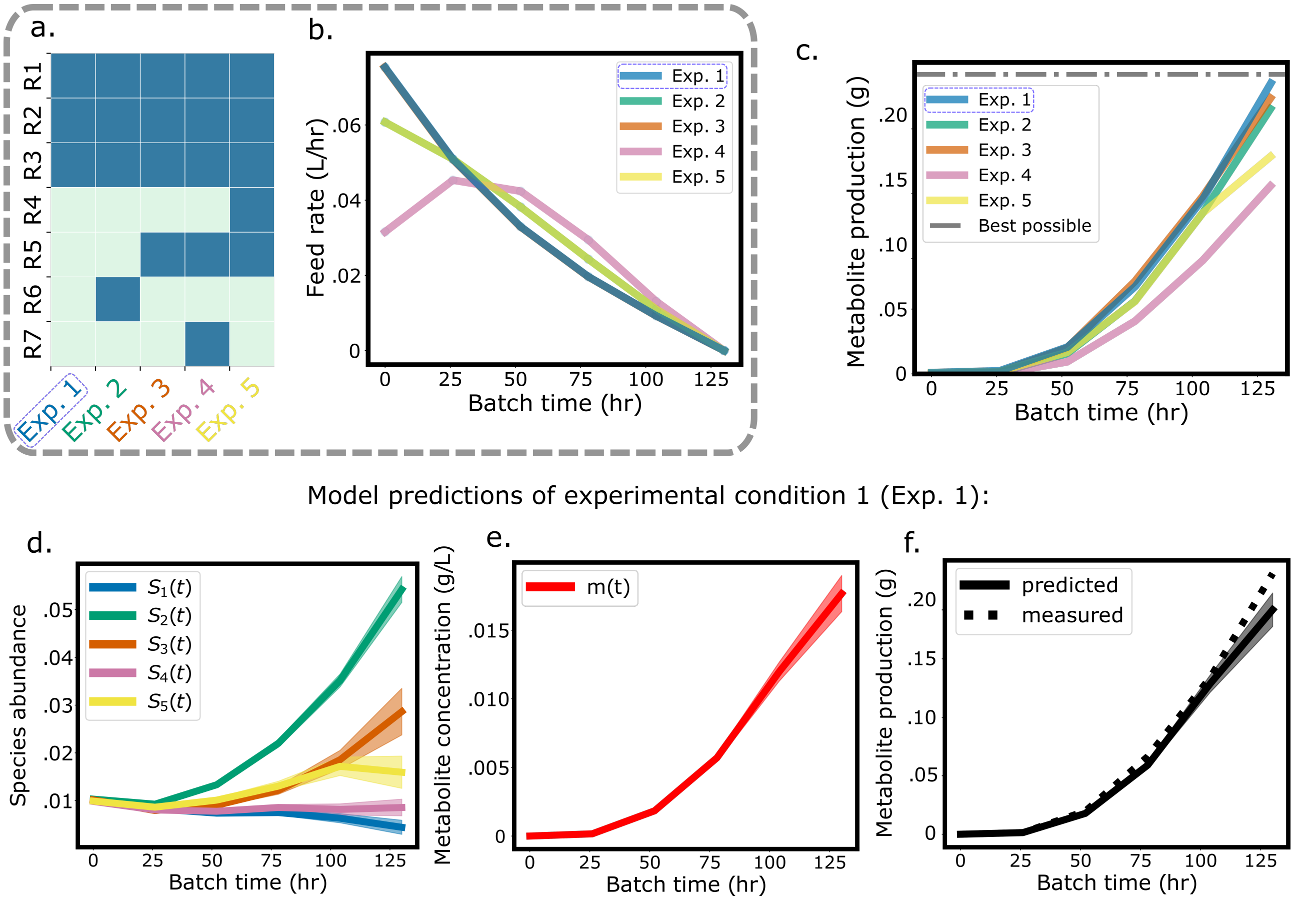


Figure 4: **Model predictions of the optimal experimental condition (Exp. 1).** (a.) The heatmap shows which resources were included in each experimental condition, where dark blue indicates the presence of a resource in the feed stream. (b.) The set of feed rates in the experimental design. (c.) Observed metabolite production in the bioreactor for each experimental condition. (d.) Experimental condition one (Exp. 1) species predictions and uncertainty intervals (mean $\pm 1$ standard deviation) (e.) Experimental condition one (Exp. 1) metabolite prediction and uncertainty interval (mean $\pm 1$ standard deviation) (f.) Prediction (mean $\pm 1$ standard deviation) of metabolite production compared to measured values.

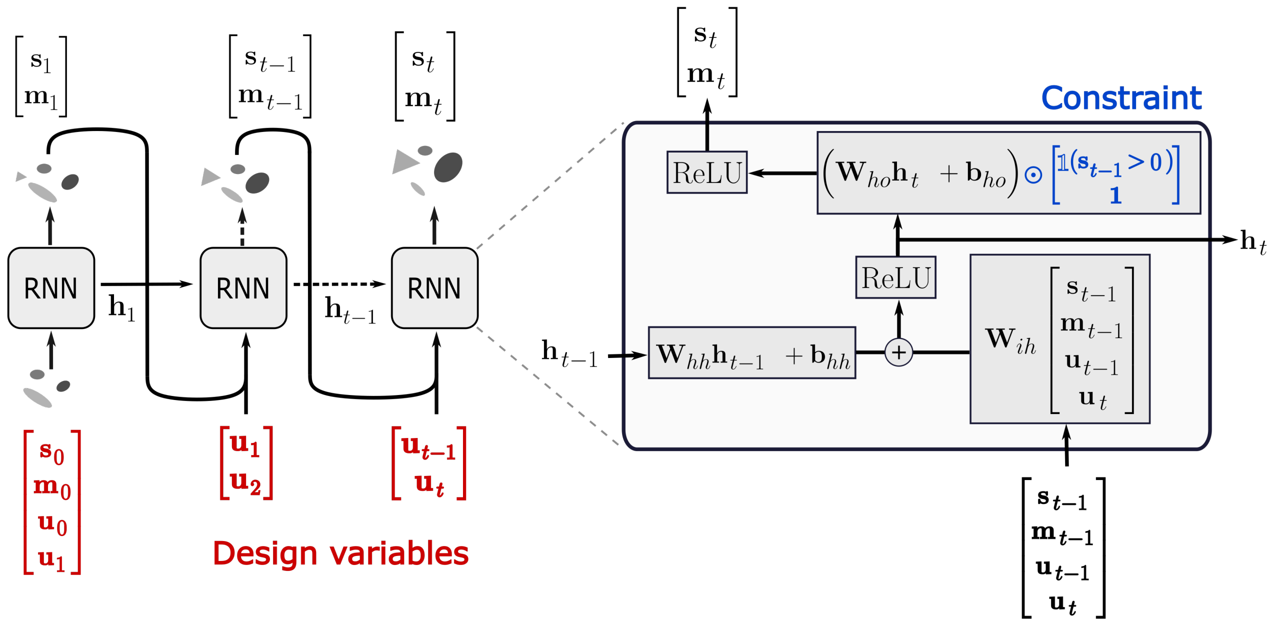


Figure 5: **Detailed view of MiRNN model architecture.** The set of model parameters of the architecture is composed of the weights and biases $\theta = \{\mathbf{W}_{hh}, \mathbf{b}_{hh}, \mathbf{W}_{ih}, \mathbf{W}_{ho}, \mathbf{b}_{ho}, \mathbf{h}_0\}$. The constraint uses an indicator function to determine whether the incoming species abundance vector, $\mathbf{s}_{t-1}$, is greater than zero. The effect of the constraint is to ensure that if a particular species is zero at time $t-1$, the model prediction of that species at time $t$ will also be zero. The ReLU function ensures that model outputs are strictly non-negative.

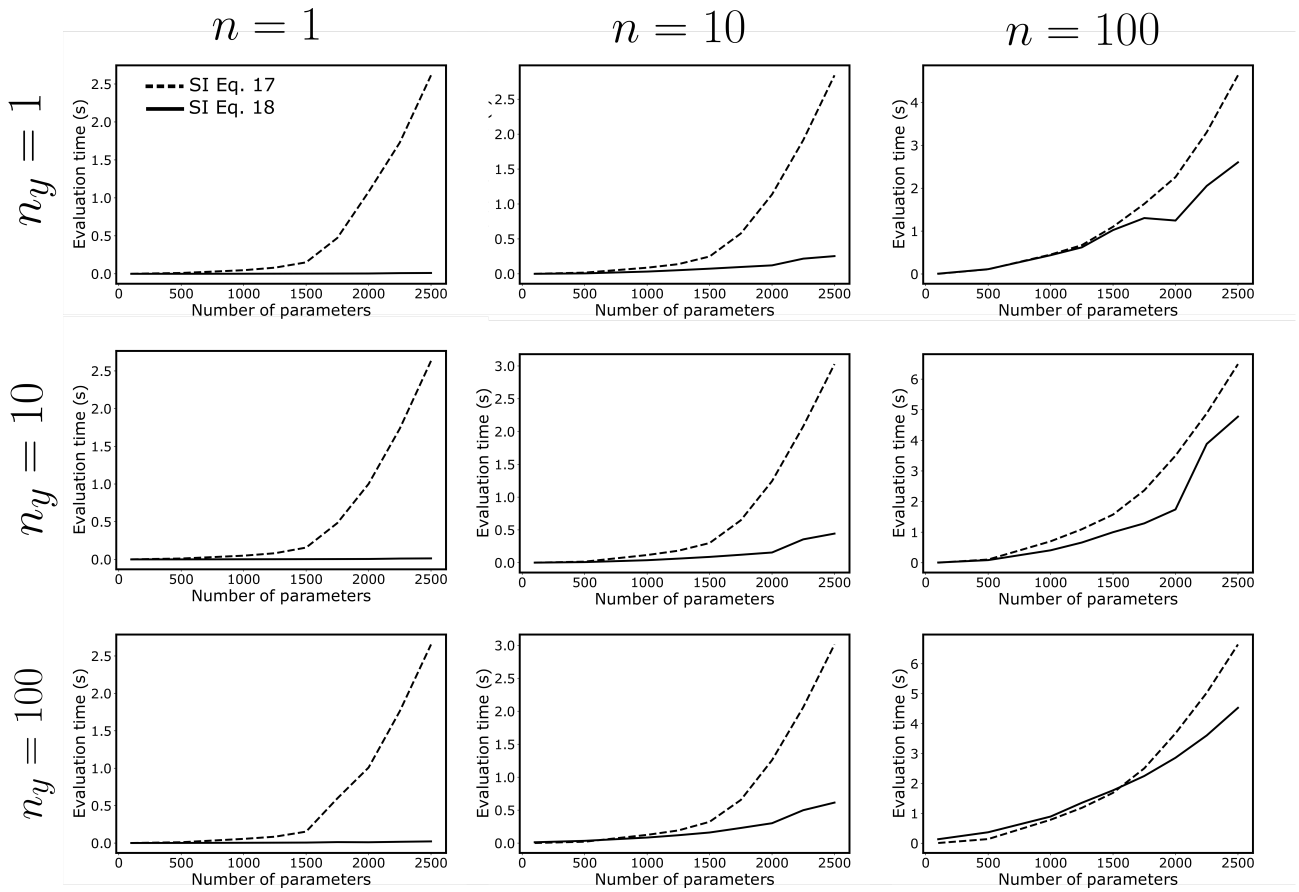


Figure 6: **Evaluation times of different methods to compute the approximate information gain.** Comparison of evaluation times of the expressions for the EIG in Eq. 18, where the number of model parameters ($n_\theta$) is varied from 0 to 2500, $n_y$ is the number of model outputs and $n$ is the number of experimental conditions in the design.

### References

[1] Christopher M Bishop and Nasser M Nasrabadi. *Pattern recognition and machine learning*. Springer, 2006.

[2] Kathryn Chaloner and Isabella Verdinelli. Bayesian experimental design: A review. *Statistical Science*, 10:273–304, 1995.

[3] F Dan Foresee and Martin T Hagan. Gauss-newton approximation to bayesian learning. In *Proceedings of international conference on neural networks (ICNN'97)*, volume 3, pages 1930–1935. IEEE, 1997.

[4] Christos Georgakis. Design of dynamic experiments: A data-driven methodology for the optimization of time-varying processes. *Industrial & Engineering Chemistry Research*, 52(35):12369–12382, 2013.

[5] Joshua E Goldford, Nanxi Lu, Djordje Bajić, Sylvie Estrela, Mikhail Tikhonov, Alicia Sanchez-Gorostiaga, Daniel Segrè, Pankaj Mehta, and Alvaro Sanchez. Emergent simplicity in microbial community assembly. *Science*, 361(6401):469–474, 2018.

[6] Juliane Liepe, Sarah Filippi, Michał Komorowski, and Michael P. H. Stumpf. Maximizing the information content of experiments in systems biology. *PLOS Computational Biology*, 9(1):1–13, 01 2013.

[7] Ali Shahmohammadi and Kimberley B McAuley. Using prior parameter knowledge in model-based design of experiments for pharmaceutical production. *AIChE Journal*, 66(11):e17021, 2020.

[8] Isabella Verdinelli and Joseph B Kadane. Bayesian designs for maximizing information and outcome. *Journal of the American Statistical Association*, 87(418):510–515, 1992.

[9] Pauli Virtanen, Ralf Gommers, Travis E. Oliphant, Matt Haberland, Tyler Reddy, David Cournapeau, Evgeni Burovski, Pearu Peterson, Warren Weckesser, Jonathan Bright, Stéfan J. van der Walt, Matthew Brett, Joshua Wilson, K. Jarrod Millman, Nikolay Mayorov, Andrew R. J. Nelson, Eric Jones, Robert Kern, Eric Larson, C J Carey, İlhan Polat, Yu Feng, Eric W. Moore, Jake VanderPlas, Denis Laxalde, Josef Perktold, Robert Cimrman, Ian Henriksen, E. A. Quintero, Charles R. Harris, Anne M. Archibald, Antônio H. Ribeiro, Fabian Pedregosa, Paul van Mulbregt, and SciPy 1.0 Contributors. SciPy 1.0: Fundamental Algorithms for Scientific Computing in Python. *Nature Methods*, 17:261–272, 2020.
